# *Citrobacter rodentium* infection disrupts colonic mucus barrier function via ablation of intercrypt goblet cells

**DOI:** 10.64898/2026.11.07.764371

**Authors:** Mahadevan Venkita Subramani, Juan S. Bailo, Alexandra Thiran, Aimee Reddy, Karl T Hansson, Gillian Blancke, Lars Vereecke, Elisabeth E.L. Nyström, Jenny Gustafsson, George M.H. Birchenough

## Abstract

The colonic mucus barrier, composed of distinct layers maintained by specialized goblet cell (GC) subtypes, forms the first line of defense against enteric pathogens. However, the individual contributions of inner (IML) and fecal (FML) mucus layers to barrier integrity, and how these layers and the GC responses are dynamically modulated during infection remain incompletely understood. Using *Citrobacter rodentium* infection in mice, we show that early colonization disrupts the inner mucus layer, increases mucus penetrability and drives alterations in faecal mucus layer thickness. Spatial imaging revealed progressive crypt infiltration and concurrent accumulation of pathogen clusters within the FML. Transcriptomic and functional analyses identified Spdef-dependent intercrypt goblet cells (icGCs) as essential for sustaining IML integrity and limiting epithelial access. Spdef-deficient mice exhibited delayed clearance and enhanced crypt invasion, highlighting the protective role of icGCs. Infection selectively depleted icGCs, collapsing the intercrypt mucus and creating permissive niches for colonization. Mechanistically, C. rodentium leveraged its type III secretion system, specifically the EspF effector to reprogram GC transcriptional responses, inducing metabolic stress and promoting apoptotic signatures in GCs specifically. These findings reveal a previously unrecognized virulence strategy in which EspF-driven GC remodeling compromises intercrypt mucus defenses, potentially facilitating pathogen persistence and transmission.

## Introduction

The intestinal epithelium constantly faces threats from luminal microbes and opportunistic pathogens. To protect this surface, the epithelium is coated with a mucus barrier primarily composed of the polymeric mucin protein Muc2, which is synthesized and secreted by intestinal goblet cells (GCs) [1]. The mucus layer provides spatial separation between the intestinal microbiota and the underlying epithelium, and its structural organization varies along the intestine. In the small intestine, the mucus layer forms a loosely organized network enriched with antimicrobial peptides (AMPs) secreted by Paneth cells and enterocytes, limiting microbial contact while supporting nutrient absorption [2, 3]. By contrast, the colon harbors a stratified mucus system, with a dense, bacteria-free inner mucus layer (IML) acting as a physical barrier against luminal threats, and a more porous outer layer that accommodates the commensals. This outer mucus layer is gradually removed by faecal matter as a result of peristalsis which contributes to the stool bound faecal mucus layer (FML) [4].

The colonic IML consists of regionally secreted substructures including crypt plume mucus that covers the region over colonic crypts and the intercrypt mucus that lies over the intercrypt regions. This highly dynamic and intricate mucus structure is maintained through ongoing mucus secretion from colonic surface GCs. GCs orchestrate this through specialized cellular programs involving Muc2 production, folding, glycosylation, transport, and exocytosis [5, 6]. Secreted polymers of Muc2 are post translationally modified by transglutaminase-mediated cross-linking. Specifically, transglutaminase 3 (Tgm3) reinforces crypt plume mucus in the distal colon by cross-linking Muc2 polymers, thereby supporting barrier integrity. Previous studies have shown that deficiency of Tgm3 leads to a more severe dextran sodium sulfate (DSS) -induced colitis and greater vulnerability to proteolytic degradation of crypt plume mucus, although its role during protection from infection remains unclear [7].

Recent insights highlight that GCs are genetically heterogenous in nature [8]. Transcriptomic profiling categorizes them into canonical and non-canonical lineages, with canonical intercrypt goblet cells (icGCs) being the most mature GCs that are functionally dependent on expression of the transcription factor SAM pointed domain containing ETS transcription factor (Spdef) [9, 10]. The icGCs produce intercrypt mucus that further reinforces IML integrity. A previous study identified Spdef-deficient mice to present spontaneous age dependent as well as DSS induced colitis due to an impaired icGC function and dysfunctional intercrypt mucus [8].

Though secretion by different GCs sustains the IML as a first line of defense, various factors including defects in secretion [11, 12], shifts in microbiota composition [13, 14], dietary influences or infection can destabilize this layer and contribute to colitis [15]. While several reports have highlighted the role of mucus in protection against enteric infections, the underlying mechanism and strategies by which pathogens disrupt this mucus barrier to colonise the intestine remain elusive. There is limited understanding of how pathogens interact with the distinct components of the mucus defense system, including the FML, IML, its subcomponents, crypt plume and intercrypt mucus, and the specialized GC subtypes responsible for their maintenance.

To delineate this, we employed *Citrobacter rodentium*, a murine pathogen that colonises the colon by overcoming the previously stated components of mucus associated protection. In murine models, Muc2 deficiency dramatically worsens *C. rodentium* infection, underlining the mucus barrier’s critical role in host defense [16]. Yet, which specific mucus sublayers and goblet-cell functions govern those very first colonisation events when the pathogen must breach the barrier, remains unknown. After crossing the IML, *C. rodentium* deploys a type III secretion system (T3SS) to tightly adhere to intestinal epithelial cells by forming attaching and effacing (A/E) lesions, a strategy prevalently employed by Enteropathogenic *E. coli* (EPEC) and Enterohaemorrhagic *E. coli* (EHEC) in humans. It has a locus of enterocyte attachment (LEE) pathogenicity island, a set of highly conserved genes that encode for bacterial effector proteins and the T3SS machinery which facilitates its attachment to the epithelium [17]. During the infection cycle *C. rodentium* causes commensal dysbiosis, triggers hyperplasia in colonic crypts, induces mucus secretion from GCs of the intestine and activates acute innate and adaptive immune responses from lamina propria residing immune cells [18–20]. However, it remains unclear whether these pathogenic mechanisms directly regulate or interact with GC subtypes, particularly in ways that compromise mucus barrier integrity or alter GC function. Understanding this interface is critical to deciphering how pathogens exploit epithelial defenses and reshape mucosal architecture during infection.

Therefore, using *C. rodentium*, we have investigated spatial and temporal changes in tissue and faecal mucus architecture that occur during infection. We demonstrate that the Spdef-dependent icGC subpopulation is a pivotal driver of mucosal defense against *C. rodentium*. In addition, our study identifies a novel and active pathogenic strategy to manipulate the expression and function of these specialized cells, thereby establishing *C. rodentium’s* colonic niche.

## Results

### C. rodentium infection disrupts the distal colonic inner mucus layer

Earlier work has shown that *C. rodentium* can closely colonize the murine colonic epithelium. In this context, the Muc2-based inner mucus layer (IML) is essential for preventing extensive mucosal injury and maintaining barrier function during infection. Beyond maintaining structural integrity, Muc2 also helps control luminal bacterial load by entrapping loosely adherent pathogens and possibly expelling them via stool shedding [11, 23]. As *C. rodentium* could disrupt the tissue-bound IML and the overlying faecal mucus layer (FML) during its infection cycle, we set out to monitor dynamic changes in these layers throughout the course of infection.

In our experiments, uninfected control mice showed gradual weight gain while infected mice did not gain or lose weight, as previously observed for wild-type (WT) C57BL/6 mice that do not normally develop systemic infections after *C. rodentium* infection (Fig. 1A). Stool-associated luminal *C. rodentium* counts peaked at approximately 10⁹ CFU/g by day 9 and remained as high as 10^6^ CFU/g by day 14 (Fig. 1B). Viable bacteria were also recovered from colonic tissues as early as 3 dpi at high titres (10⁷ CFU/g tissue), peaking at 7dpi (∼ 10^8^ CFU/g tissue), which remained high (∼10⁶ CFU/g tissue) even at 14 dpi, confirming robust mucosal colonisation (Fig. 1C). Collectively, these data confirm the dynamic trajectory of *C. rodentium* infection, marked by progressive faecal bacterial shedding alongside sustained mucosal colonization.

**Figure 1:**
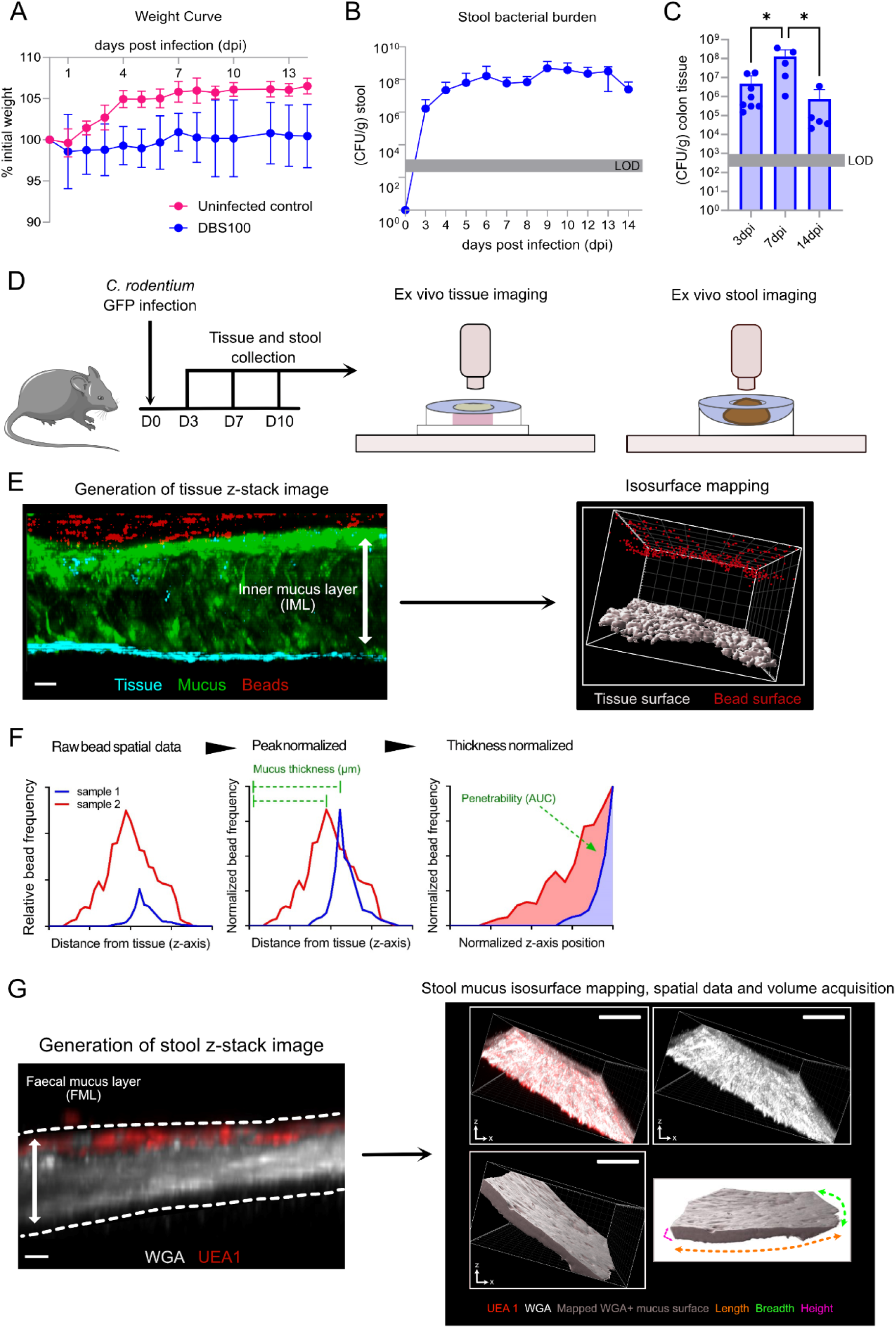
Experimental model of C. rodentium infection and ex vivo assessment of the colonic mucus barrier. **A)** Body weight changes in uninfected control (pink) and C. rodentium DBS100-infected (blue) mice over 14 days post-infection (dpi), expressed as a percentage of initial body weight. **B)** Stool C. rodentium burden quantified as colony-forming units per gram of stool (CFU/g) at different time points post-infection. The grey shaded area indicates the limit of detection (LOD). **C)** Distal colonic tissue C. rodentium burden quantified as CFU/g at 3, 7 and 14 dpi. The grey shaded area indicates the LOD. **D)** Schematic of the experimental design. Mice were orally infected with GFP-expressing C. rodentium DBS100 at day 0, followed by tissue and stool collection at 3, 7 and 10 dpi for ex vivo confocal imaging. **E)** Representative confocal z-stack of distal colonic tissue showing the tissue surface (cyan), mucus layer (green) and fluorescent microbeads (red). The white arrow indicates the inner mucus layer (IML). Three-dimensional isosurface reconstruction of the tissue surface (white) and microbeads (red) was used to determine their relative z-axis positions. **F)** Schematic of mucus penetrability analysis based on microbead spatial distribution. Graphs show raw microbead frequency distributions (left), peak-normalized frequency distributions aligned to the mucus surface (middle), and thickness-normalized distributions (right). The shaded area under the normalized curves represents microbead penetrability, quantified as area under the curve (AUC). **G)** Representative confocal z-stack of a stool pellet showing the faecal mucus layer (FML) stained with WGA (white) and UEA-1 (red). Dashed lines indicate the boundaries of the FML. Three-dimensional isosurface reconstructions were used to map the stool-associated mucus layer and determine its volume and dimensions (length, breadth and height). Data represent n = 6-10 mice per group, where indicated. Body weight and stool bacterial burden are presented as mean ± SEM (A, B), whereas tissue bacterial burden is presented as median and interquartile range with individual data points (C). Statistical comparisons were performed using two-way ANOVA followed by Fisher’s LSD test (A) or one-way ANOVA followed by Fisher’s LSD test (C). Statistical significance is indicated as p < 0.05 (*) and p < 0.01 (**). Image scale bars are 50 µm (E) and 100 µm (G).

The pronounced mucosal bacterial burden confirmed that *C. rodentium* can breach the mucus barrier to establish an epithelial niche, followed by dissemination via faecal shedding. We hypothesised that the pathogen modulates biophysical properties of the colonic mucus layer, specifically its thickness and penetrability to facilitate this process. To explore this, we quantified both the inner mucus layer (IML), which adheres closely to the epithelium, and the stool-associated faecal mucus layer (FML). IML characteristics were assessed using an ex *vivo* mucus penetrability assay in combination with confocal imaging using bacteria sized beads, as previously described [24]. Fresh stool pellets were placed in central indentations of tac-clay domes on custom slides to ensure they remained intact and immobile.

A chamber was placed on top of the mounted stool to isolate a portion of the stool pellet surface. This apical surface was stained and imaged via confocal microscopy and processed through our analysis pipeline to quantify FML thickness (Fig. 1D–G). Our analyses revealed a progressive, infection-dependent thinning of the IML, alongside a significant increase in *ex vivo* mucus penetrability at 7 and 10 dpi (Fig. 2A–D). Given the observed IML thinning and increased penetrability, we next examined whether *C. rodentium* infection altered mucus growth kinetics. In live distal colonic tissue, baseline IML growth over 30 min remained unchanged between uninfected control mice and those at 3 dpi but fell significantly at 7 and 10 dpi compared to controls (Fig. 2E), indicating impaired replenishment of the inner layer during *C. rodentium* infection.

**Figure 2:**
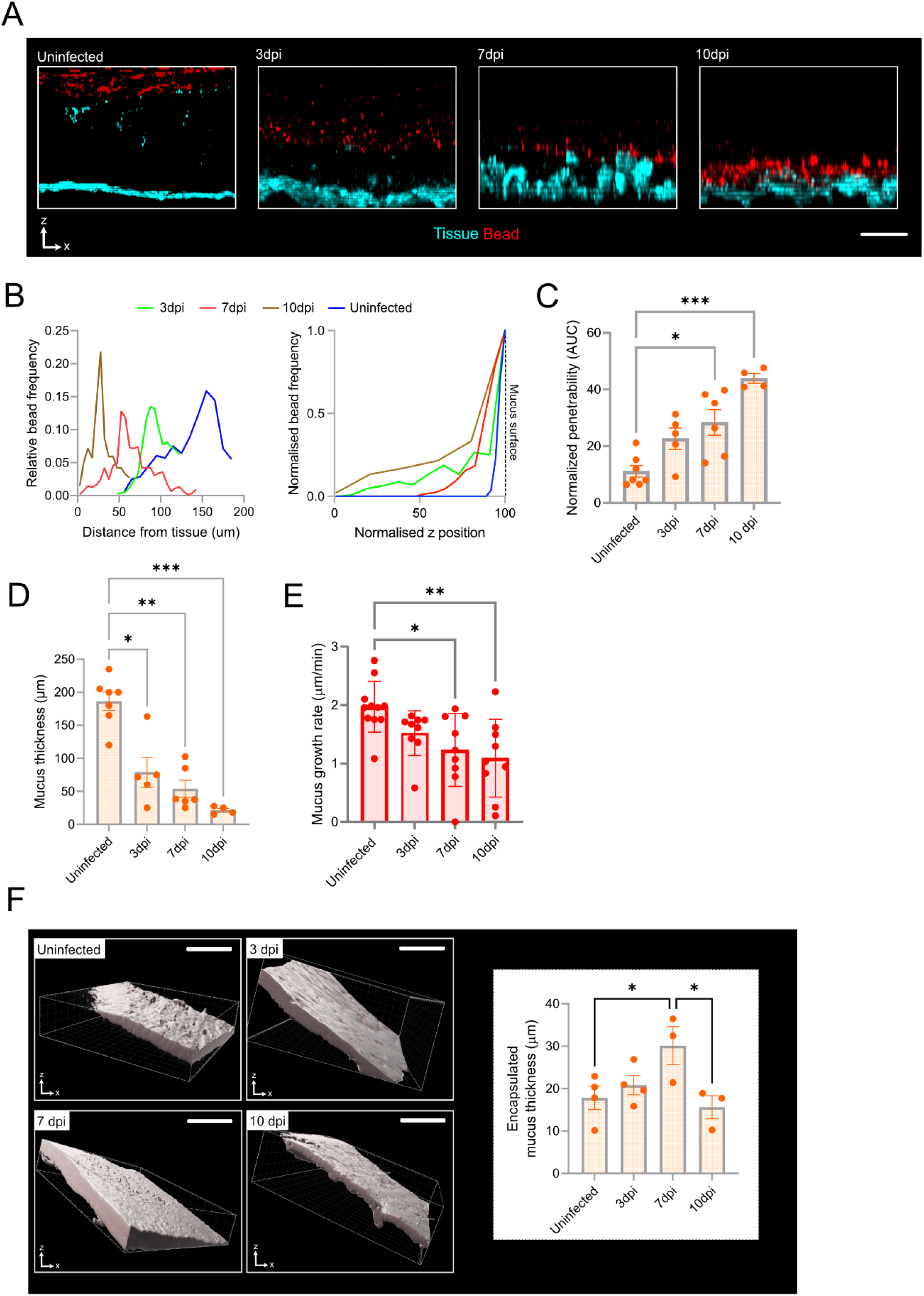
*C. rodentium* infection disrupts the distal colonic inner mucus layer barrier. **A)** Representative confocal z-stacks of distal colonic tissue from uninfected and *C. rodentium*-infected mice at 3, 7 and 10 days post-infection (dpi). Images show x/z-axis cross-sections of the tissue surface (cyan) overlaid with fluorescent microbeads (red), illustrating changes in mucus barrier penetrability during infection. **B)** Microbead spatial distributions across the mucus layer at different infection time points. Graphs show relative microbead frequency plotted against distance from the tissue surface (left) and normalized microbead frequency plotted against normalized z-axis position (right), with the mucus surface aligned to the position of maximum microbead accumulation. **C)** Mucus barrier function quantified by normalized microbead penetrability, expressed as area under the curve (AUC), in uninfected and infected mice at 3, 7 and 10 dpi. **D)** Inner mucus layer (IML) thickness quantified by ex vivo mucus thickness measurements in distal colonic tissue from uninfected and infected mice at 3, 7 and 10 dpi. **E)** IML growth rate quantified by ex vivo mucus growth measurements in distal colonic tissue from uninfected and infected mice at 3, 7 and 10 dpi. **F)** Representative three-dimensional isosurface reconstructions of the faecal mucus layer (FML) from stool samples of uninfected and infected mice at 3, 7 and 10 dpi (left). Encapsulated mucus thickness quantified from the reconstructed FML at each infection time point (right). Data represent n = 4-7 mice per group (A–D), n = 8-10 mice per group (E), or n = 3-4 mice per group (F). Bar graphs show mean ± SEM with individual data points overlaid (C–F). Statistical comparisons were performed using the Kruskal–Wallis test followed by Dunn’s multiple comparisons test (C) or one-way ANOVA followed by Fisher’s LSD post hoc test (D–F). Statistical significance is indicated as p < 0.05 (\**), p < 0.01 (**), and p < 0.001 (\*\*\**). Image scale bars represent 75 µm (A) and 100 µm (F).

Despite a reduction in distal colonic baseline mucus replenishment, total expelled mucus accumulated along the proximal-distal axis, as reflected by the fact that FML thickness did not decrease. Instead, FML thickness rose markedly up to 7 dpi (Fig. 2F), suggesting a compensatory secretory response from regions proximal to distal colon. By 10 dpi, FML thickness began to decline, yet remained comparable to uninfected levels (Fig. 2F).

Together, these findings indicate that *C. rodentium* colonises the colonic epithelium and the infection subsequently drives modulation of the thickness and barrier quality of the IML. Interestingly, FML thickness increases during early infection, which could reflect pathogen-driven mucus secretion or a host effort to flush out bacteria. Since distal colonic mucus growth rate deteriorates from 7 dpi onwards, this rise in FML thickness likely reflects more proximal rather than local colonic mucus secretion.

### Mucus barrier disruption correlates with C. rodentium crypt infiltration and faecal mucus entrapment

Given the observed reduction in both IML barrier function and thickness during *C. rodentium* infection, coupled with a sustained high mucosal bacterial burden, we hypothesised that the pathogen exploits this compromised barrier to access and colonise the colonic crypts. To investigate this, we employed an ex vivo tissue imaging assay using fluorescent UEA-1 (intercrypt mucus) and WGA (crypt mucus and epithelial glycocalyx) lectin-based imaging of live colonic tissue collected from mice infected with GFP-expressing *C. rodentium [8]*.

Direct adherence of GFP⁺ *C. rodentium* to the epithelial surface was evident as early as 3 days post-infection (dpi), with progressive infiltration into the crypts observed at 7 and 10 dpi (Fig. 3A). This spatial progression correlated with a notable reduction in mucus coverage, as visualised by lectin staining (Fig. 3A) and supporting our previous barrier function data (Fig. 2). Moreover, crypt-localised colonisation appeared to be associated with disruption of overall mucosal epithelial integrity, with increasing tissue deterioration apparent by WGA staining at 7 and 10 dpi (Fig. 3A). This was associated with a relatively stable mean fluorescence intensity (MFI) of GFP+ cells across 3–10 dpi, without a clear upward trend (Fig. 3F).

**Figure 3:**
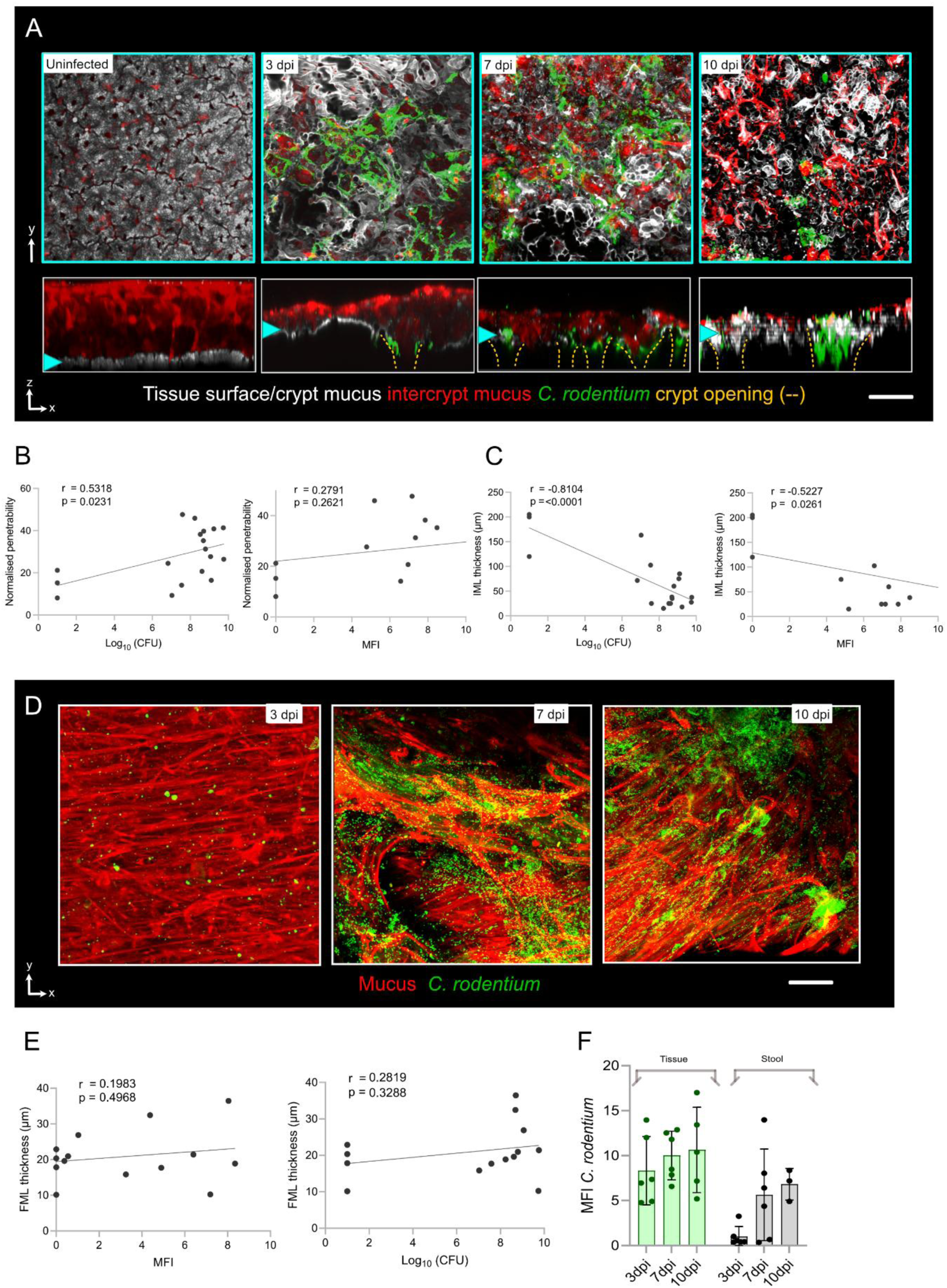
*C. rodentium* infection is associated with mucus barrier disruption, crypt colonization and faecal mucus entrapment. **A)** Representative confocal micrographs of distal colonic tissue from uninfected and *C. rodentium*-infected mice at 3, 7 and 10 days post-infection (dpi). Images show x/y-axis views (top) and x/z-axis cross-sections (bottom) of the tissue surface. Yellow dashed lines indicate crypt openings, and cyan arrowheads indicate the tissue surface. **B)** Correlation between normalized mucus penetrability and luminal *C. rodentium* burden, quantified as log₁₀ CFU (left), or tissue-associated bacterial GFP mean fluorescence intensity (MFI; right). Linear regression lines and Pearson correlation coefficients (r) and p-values are shown. **C)** Correlation between inner mucus layer (IML) thickness and luminal *C. rodentium* burden, quantified as log₁₀ CFU (left), or tissue-associated bacterial GFP MFI (right). Linear regression lines and Pearson correlation coefficients (r) and p-values are shown. **D)** Representative confocal micrographs of stool-associated faecal mucus layer (FML) from *C. rodentium*-infected mice at 3, 7 and 10 dpi. Images show mucus stained with UEA-1 (red) and GFP-expressing *C. rodentium* (green). **E)** Correlation between FML thickness and stool-associated *C. rodentium* burden, quantified as GFP MFI (left) or log₁₀ CFU (right). Linear regression lines and Pearson correlation coefficients (r) and p-values are shown. **F)** Tissue-associated (green) and stool-associated (grey) *C. rodentium* GFP MFI quantified at 3, 7 and 10 dpi. Data are presented as individual observations with linear regression lines (B, C, E) or mean ± SEM with individual data points overlaid (F). Correlations were assessed using Pearson’s correlation analysis (B, C, E), with corresponding r and p-values indicated in each graph. Statistical comparisons were performed using one-way ANOVA followed by Fisher’s LSD post hoc test (F). Statistical significance is indicated as p < 0.05 (*), p < 0.01 (), and p < 0.001 (*). Image scale bars represent 50 µm (A, D).

To further interrogate the relationship between mucus barrier integrity and bacterial colonization, we performed correlation analysis. Notably, normalized mucus penetrability and IML thickness values consistently correlated significantly with luminal *C. rodentium* burden, whereas their correlations with tissue-associated bacterial MFI were either weak or not significant (Fig. 3B, C). These data are more consistent with a model in which epithelial colonization precedes and drives IML disruption, leading to mucus layer loss and increased shedding into the lumen, rather than barrier breakdown enabling additional colonization at the same sites (Fig. 3B, C)

Previously we observed an increase in FML thickness along the course of the infection which we hypothesized to be linked with an increase in live *C. rodentium* in the feces. To assess this, we visualised and quantified the presence of *C. rodentium* within the stool-associated mucus. At 3 dpi, small microcolonies were detected within the faecal mucus layer (FML) (Fig. 3D). By 7 and 10 dpi, these expanded into larger bacterial clusters that co-localized with the mucus layer, signifying a potential role for mucus in pathogen entrapment and shedding (Fig. 3D). Notably, bacteria were predominantly localized on the outer surface of the FML, indicating a pattern of shedding from the distal epithelium. Supporting this, median fluorescence intensity (MFI) analysis revealed a gradual accumulation of GFP⁺ *C. rodentium* signal in the stool, peaking at 7 dpi (Fig. 3E, F)

Together, the data indicate that epithelial colonization drives subsequent IML disruption. Tissue-associated GFP signal remains broadly stable, while mucus penetrability and IML measures correlate with luminal burden, pointing to increased shedding rather than deeper tissue expansion. Additionally, stool-associated bacterial clusters and stool GFP peak at 7 dpi, consistent with barrier damage suggesting a route for luminal release and further transmissibility.

### C. rodentium infection modulates colonic GC abundance, survival and proliferative capacity of GCs

Given the spatial disruption of mucus architecture observed during *C. rodentium* infection and its influence on colonisation kinetics, we hypothesized that these changes could be potentially influenced by underlying alterations to GC dynamics driven by infection. To test this, we utilised our reporter RedMUC2^98trTg^ strain, which expresses an mCherry-tagged Muc2 reporter, and employed a flow cytometry approach to evaluate viable GC frequency, Muc2 expression (via mCherry intensity), and proliferative capacity (via Ki67 labelling) across infection stages (Fig. 4A-B). Flow cytometry on isolated colonic epithelial cells was used after sequential gating to exclude doublets/dead cells, select EpCAM⁺ cells, gate mCherry-Muc2⁺ goblet cells, and identify Ki67⁺ proliferative subsets (Fig. 4B). RedMUC2^98trTg^ infected with *C. rodentium* exhibited 5% body weight loss by 6 days post-infection (dpi), alongside a progressive increase in stool bacterial burden peaking at 10^8^ CFU/g by 10 dpi (Fig. 5A–B). These trends paralleled the infection dynamics previously seen in our wild-type animals.

**Figure 4:**
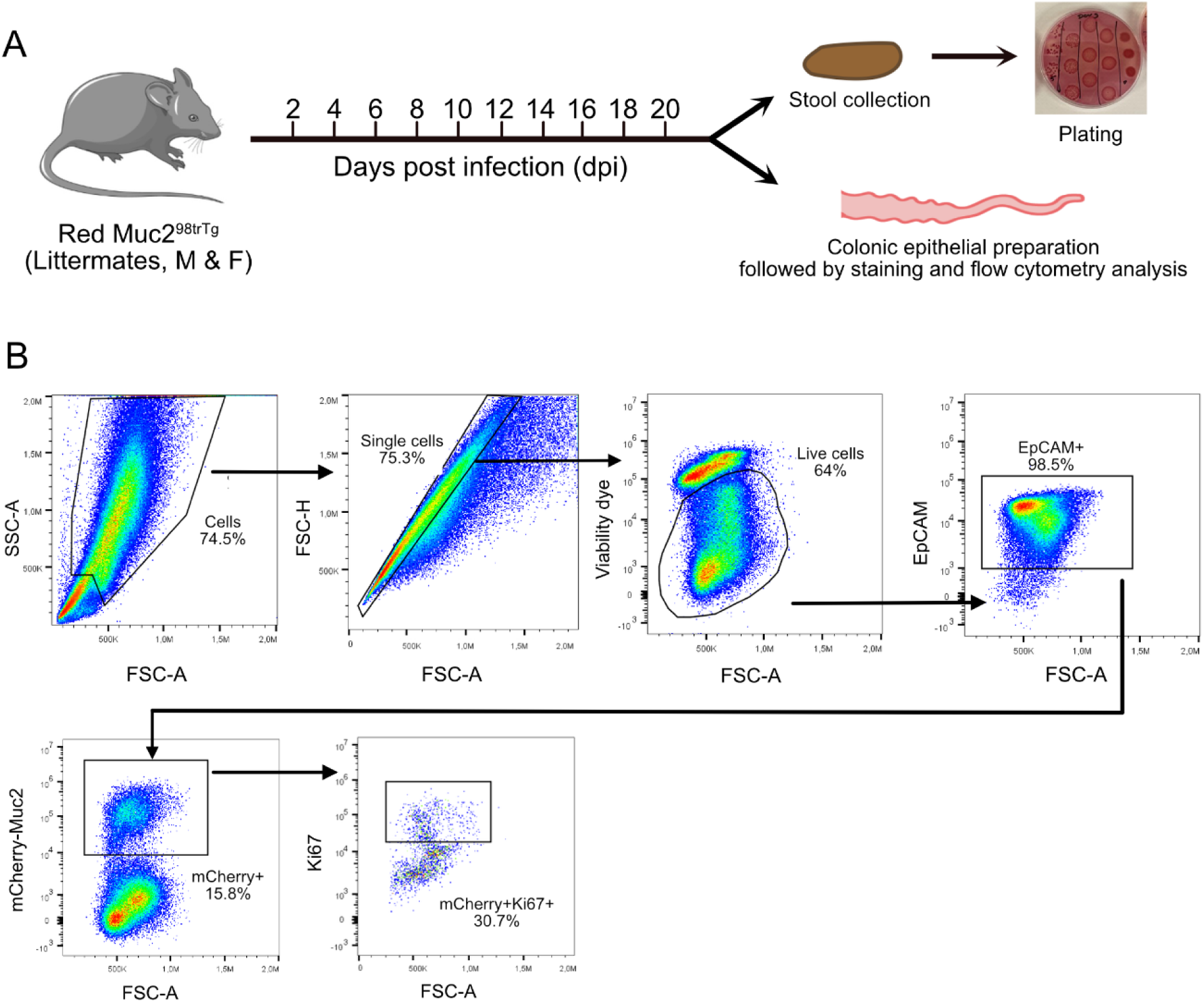
Experimental design and flow cytometry gating strategy for colonic epithelial cell analysis. **A)** Schematic of the experimental design using Red Muc2^98trTg mice of both sexes from different litters. Stool samples were collected at indicated days post-infection (dpi) for bacterial burden quantification by plating. Colonic epithelial cells were isolated for flow cytometric analysis of Muc2-expressing goblet cells (mCherry-Muc2) and proliferating epithelial cells (Ki67). **B)** Representative flow cytometry plots showing the sequential gating strategy for colonic epithelial cell populations. Cells were identified based on forward scatter area (FSC-A) and side scatter area (SSC-A), followed by single-cell selection using FSC-A and forward scatter height (FSC-H). Live cells were identified by exclusion of the viability dye, followed by selection of EpCAM-positive epithelial cells. Muc2-expressing goblet cells were identified by mCherry fluorescence, and Ki67-positive cells were subsequently quantified within the mCherry-positive population. Representative percentages of gated populations are indicated in each plot.

**Figure 5:**
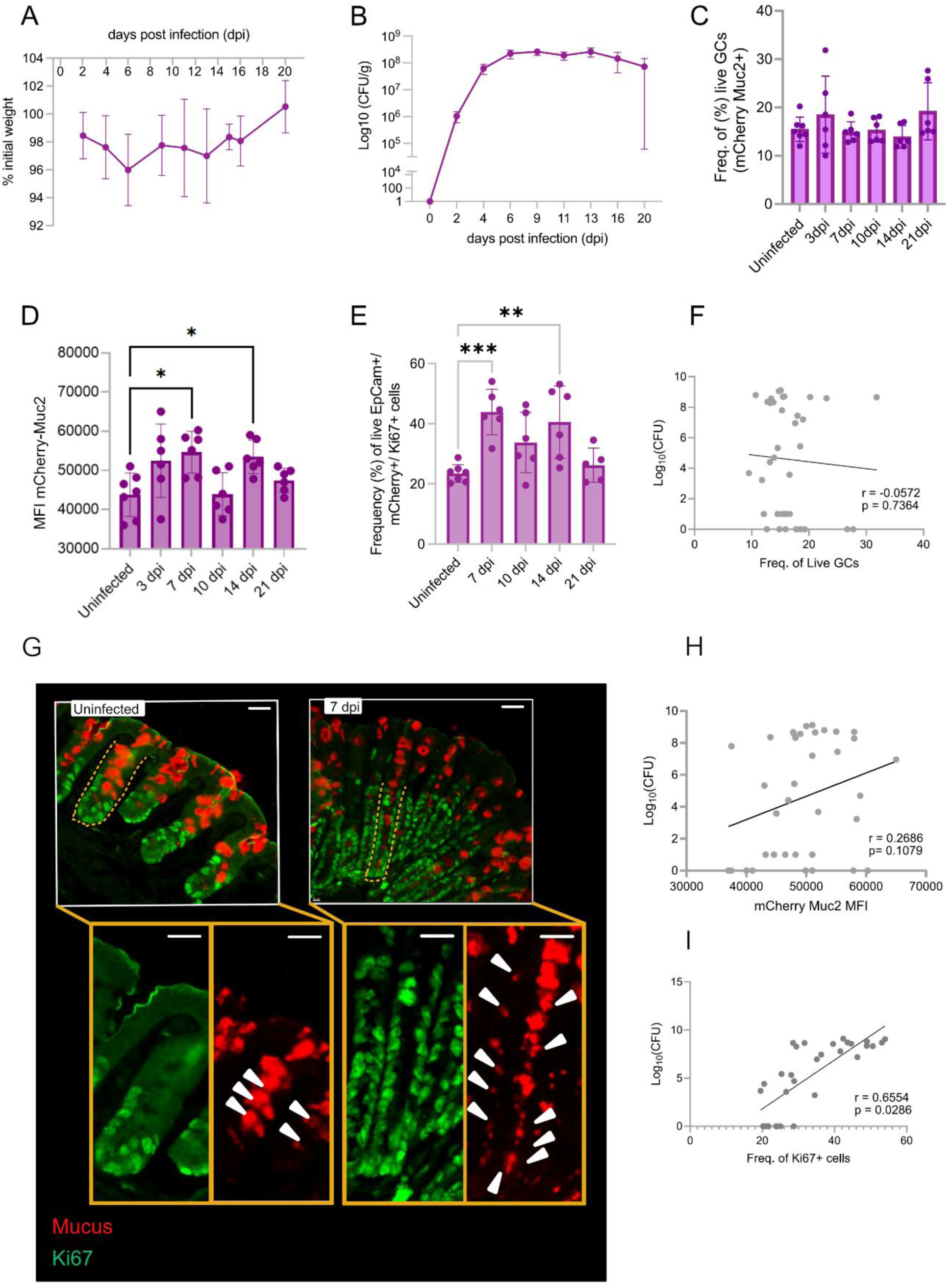
C. rodentium infection modulates colonic goblet cell proliferation and Muc2 expression. **A)** Body weight changes in C. rodentium-infected RedMuc298trTg mice over 20 days post-infection (dpi), expressed as a percentage of initial body weight. **B)** Stool C. rodentium burden quantified as colony-forming units per gram of stool (CFU/g) over 20 dpi. **C)** Frequency of live Muc2-expressing GCs, identified as EpCAM⁺ mCherry⁺ cells, in distal colonic epithelial cell populations from uninfected and infected mice at 3, 7, 10, 14 and 21 dpi. **D)** Muc2 expression quantified by mCherry mean fluorescence intensity (MFI) in EpCAM⁺ mCherry⁺ goblet cells at different infection time points. **E)** Frequency of proliferating GCs, identified as Ki67⁺ cells within the live EpCAM⁺ mCherry⁺ population, in uninfected and infected mice at 7, 10, 14 and 21 dpi. **F)** Correlation between the frequency of live GCs and stool C. rodentium burden, quantified as log₁₀ CFU. Linear regression line and Pearson correlation coefficient (r) and p-value are shown. **G)** Representative confocal micrographs of distal colonic crypts from uninfected and C. rodentium-infected mice at 7 dpi. Yellow dashed lines outline selected crypt regions, and white arrowheads indicate Muc2⁺Ki67⁺ double-positive cells. Enlarged images of the selected crypt regions are shown below. **H)** Correlation between mCherry-Muc2 MFI and stool C. rodentium burden, quantified as log₁₀ CFU. Linear regression line and Pearson correlation coefficient (r) and p-value are shown. **I)** Correlation between the frequency of Ki67-positive goblet cells and stool C. rodentium burden, quantified as log₁₀ CFU. Linear regression line and Pearson correlation coefficient (r) and p-value are shown. Data represent n = 5-7 mice per group, where indicated. Body weight and stool bacterial burden are presented as mean ± SEM (A, B), whereas bar graphs show mean ± SEM with individual data points overlaid (C–E). Statistical comparisons were performed using one-way ANOVA followed by Fisher’s LSD post hoc test (D, E). Correlations were assessed using Pearson’s correlation analysis (F, H, I), with corresponding r and p-values indicated in each graph. Statistical significance is indicated as p < 0.05 (*), p < 0.01 (**), and p < 0.001 (***). Image scale bars represent 50 µm (G).

Our analysis of live colonic epithelial cells (EpCAM⁺) revealed a steady unaltered proportion of viable GCs (Live/Dead⁻ mCherry+) throughout the infection with an average of 18% of total live cells obtained (Fig. 5C). However, mCherry–Muc2 MFI rose by ∼30% from baseline, peaking at 7 dpi, then returned to pre-infection levels by 21 dpi (Fig. 5D). Interestingly, the frequency of proliferative GCs (EpCAM⁺ mCherry⁺ Ki67⁺) increased from 20% at uninfected levels to 40% at 7 dpi, before returning to 20% at 21 dpi (Fig. 5E). To visually map the region of expansion within the crypts, we performed immunostaining of colonic sections with the proliferation marker Ki67. We confirmed upward crypt expansion of Ki67⁺ GCs at 7 dpi, consistent with crypt hyperplasia, a hallmark of *C. rodentium* infection (Fig. 5G) [20, 23].

Furthermore, we performed correlation analysis between %live GCs, mCherry MFI, %Ki67+GCs and luminal bacterial burden. We found that neither live GCs nor mCherry MFI had a significant correlation with luminal bacterial burden in these animals (Fig. 5F & H). However, the bacterial load in stool of these mice positively correlated with the frequency of proliferative GCs across time points (Fig. 5I), suggesting that luminal pathogen burden may directly influence GC proliferation in response to infection.

In summary, *C. rodentium* infection in RedMUC2^98trTg^ elicits a dynamic GC response marked by moderate increase in mCherry MFI, and a pathogen dependent enhancement of GC proliferation. However, as we observed a stable pattern of GC frequency throughout the infection, our results suggest that the previously observed IML phenotypes occur independently of GC loss or reduced Muc2 expression.

### Spdef regulates C. rodentium infection and clearance

Structural integrity of IML is regulated by two spatially distinct GC populations: crypt-resident GCs that secrete plume mucus to protect crypt openings, and intercrypt GCs that generate the intercrypt mucus network as a substructure of the IML. The fact that *C. rodentium* colonization is followed by IML breakdown implies that the pathogen may directly undermine these discrete mucus substructures, compromising their collective ability to uphold the IML’s barrier function.

Therefore, to identify which factors regulating IML barrier function are affected during infection, we performed bulk RNA sequencing of sorted colonic GCs at 10 days post-infection and investigated expression of genes that may contribute to mucosal protection. Notably, we observed a total of 423 upregulated and 467 downregulated genes that were significantly altered in GCs during infection (Fig. 6A).

**Figure 6:**
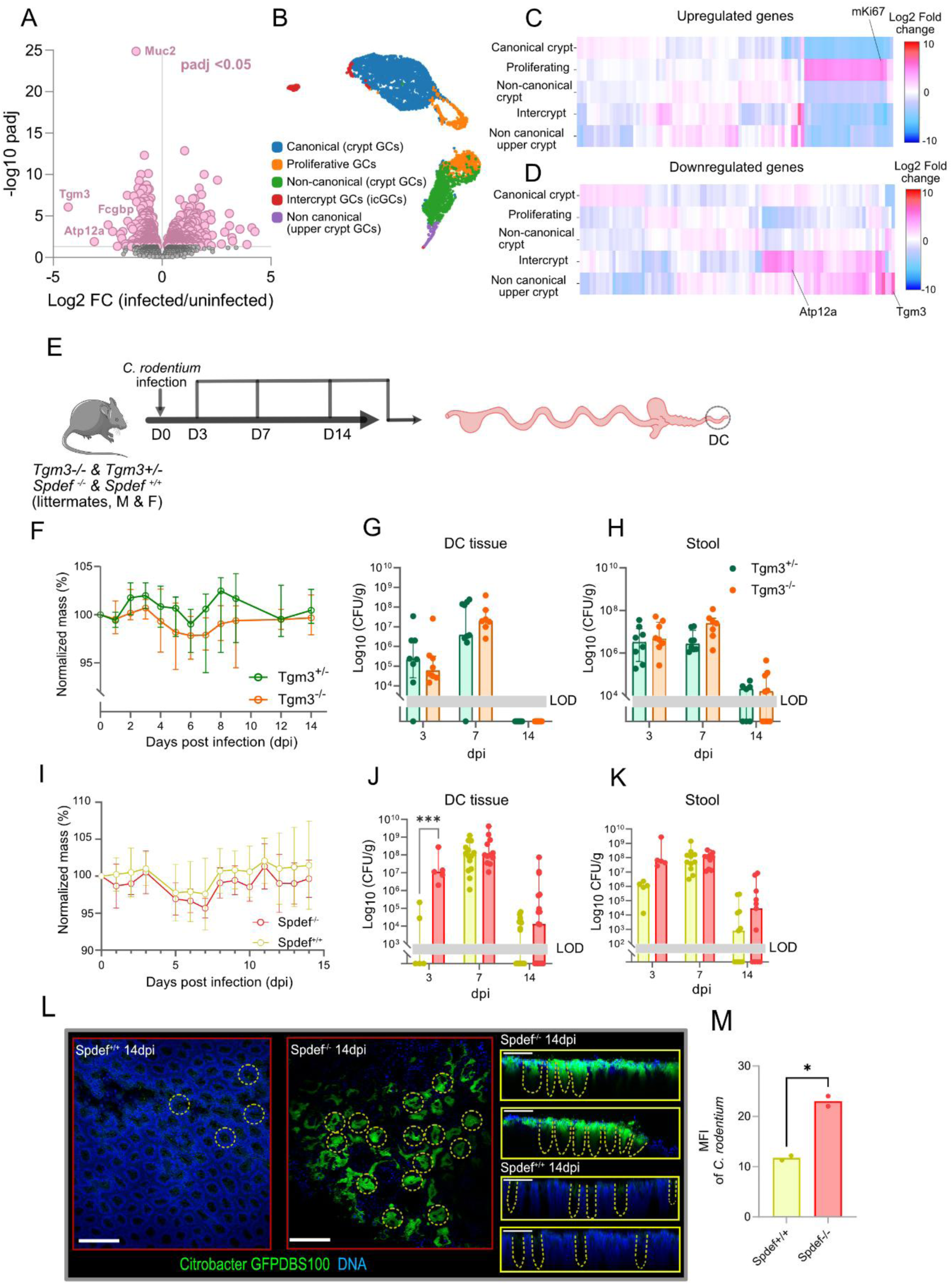
Spdef regulates C. rodentium infection and clearance. **A)** Volcano plot of differentially expressed genes from bulk RNA sequencing of sorted colonic goblet cells from Muc2-YFP reporter mice at 10 days post-infection (dpi). The x-axis indicates log₂ fold change (infected/uninfected), and the y-axis represents −log₁₀ adjusted p-value (padj). Differentially expressed genes with padj < 0.05 are highlighted in pink, with selected genes indicated. **B)** Uniform Manifold Approximation and Projection (UMAP) of single-cell transcriptomic data showing five GC sub-populations: **C)** Heatmap showing the expression patterns of upregulated genes across the five GC populations. Colour scale represents log₂ fold change, with selected genes indicated (C & D). **D)** Heatmap showing the expression patterns of downregulated genes across the same GC populations. **E)** Schematic of the experimental design using Tgm3⁺/⁻ and Tgm3⁻/⁻ or Spdef⁺/⁺ and Spdef⁻/⁻ littermate mice of both sexes. Mice were infected with C. rodentium at day 0, followed by tissue and stool collection at 3, 7 and 14 dpi. **F)** Body weight changes in C. rodentium-infected Tgm3⁺/⁻ (green) and Tgm3⁻/⁻ (orange) mice over 14 dpi, expressed as a percentage of initial body weight. **G)** Distal colonic tissue C. rodentium burden quantified as colony-forming units per gram of tissue (CFU/g) in Tgm3⁺/⁻ and Tgm3⁻/⁻ mice at 3, 7 and 14 dpi. **H)** Stool C. rodentium burden quantified as CFU/g in Tgm3⁺/⁻ and Tgm3⁻/⁻ mice at 3, 7 and 14 dpi. **I)** Body weight changes in C. rodentium-infected Spdef⁺/⁺ (yellow) and Spdef⁻/⁻ (red) mice over 14 dpi, expressed as a percentage of initial body weight. **J)** Distal colonic tissue C. rodentium burden quantified as CFU/g in Spdef⁺/⁺ and Spdef⁻/⁻ mice at 3, 7 and 14 dpi. **K)** Stool C. rodentium burden quantified as CFU/g in Spdef⁺/⁺ and Spdef⁻/⁻ mice at 3, 7 and 14 dpi. **L)** Representative confocal micrographs of distal colonic whole mounts from Spdef⁺/⁺ and Spdef⁻/⁻ mice at 14 dpi, showing GFP-expressing C. rodentium (green) and DNA (blue). Yellow dashed lines indicate crypt openings. Images on the right show x/z-axis cross-sections illustrating bacterial localization within colonic crypts. **M)** Quantification of C. rodentium GFP mean fluorescence intensity (MFI) in distal colonic tissue from Spdef⁺/⁺ and Spdef⁻/⁻ mice at 14 dpi. Data represent n = 7-8 mice per genotype (F-H), n = 5-14 mice per genotype (I-K), or n = 2 mice per genotype (L, M). Body weight changes are presented as mean ± SEM (F, I), whereas bacterial burden and fluorescence intensity measurements are presented as mean ± SEM with individual data points overlaid (G, H, J, K, M). Statistical comparisons were performed using ANOVA followed by Fisher’s LSD post hoc test (J) or the Mann-Whitney U test (M). Statistical significance is indicated as p < 0.05 (*), p < 0.01 (**), and p < 0.001 (***). Grey shaded areas indicate the limit of detection (LOD) in bacterial burden measurements (G, H, J, K). Image scale bars represent 50 µm (L).

Recent single cell transcriptomic data has identified gene expression signatures of GC subpopulations that are defined by expression of canonical or non-canonical markers and their spatial position along the crypt-intercrypt epithelial axis. We therefore mapped expression of genes differentially regulated by *C. rodentium* infection onto five GC clusters: canonical crypt GCs, proliferative GCs, non-canonical crypt GCs, intercrypt GCs (icGCs), and non-canonical upper crypt GCs, to infer GC subtype-specific alterations during infection (Fig. 6B) [8]. When we compared the expression of all genes upregulated and downregulated by infection within these clusters, we found that upregulated genes (including *mKi67*) were associated with enhanced expression in the proliferative GC cluster, while down regulated genes such as *Tgm3* and *Atp12a* mapped onto the non-canonical upper crypt GC and icGC clusters respectively (Fig. 6C-D). Together, these findings indicated that as well as inducing GC proliferation, infection downregulated functionally relevant genes associated with upper crypt and icGC populations,

We first investigated the potential role of Tgm3 downregulation as prior work demonstrated that this protein is involved in stabilizing the IML *via* cross-linking of Muc2 in crypt plume mucus [7]. We therefore infected both *Tgm3^+/-^* and *Tgm3^-/-^* littermates with *C. rodentium* and monitored infection kinetics (Fig 6E). Both genotypes exhibited comparable weight loss (∼5–6% by 6 dpi; Fig. 6F). Assessment of bacterial burden in the distal colon revealed no significant differences at 3, 7, or 14 dpi, with both genotypes reaching peak stool CFU levels of ∼10^7^ CFU/g and mucosal burdens of ∼10^8^ CFU/g by 7 dpi (Fig. 6G, H). These results suggest that Tgm3-mediated cross-linking of crypt plume mucus is not a critical determinant of protection against this pathogen.

As multiple markers of icGCs were also downregulated by *C. rodentium* infection we also investigated the role of this population in protection. To achieve this, we made use of the transcription factor Spdef, as it is essential for functioning of icGCs [9, 10]. We infected both *Spdef^+/+^* and *Spdef^-/-^* littermated animals with *C. rodentium.* By contrast to *Tgm3^-/-^* infection, *Spdef^-/-^* mice showed markedly impaired control of infection. *Spdef^-/-^* mice experienced a slight weight loss (not significant) compared to *Spdef^+/+^* control animals (Fig. 6I). However, at 3 dpi, colonic tissue CFU counts in *Spdef^-/-^*mice were ∼2 log higher than in *Spdef^+/+^* controls (10^6^ vs 10^4^ CFU/g), and stool burdens were similarly elevated, though not significant (Fig. 6J, K). Although bacterial loads converged by 7 dpi, *Spdef^-/-^* animals showed a delayed tendency towards clearance, maintaining elevated tissue CFUs at 14 dpi (10^6^ vs 10⁴ CFU/g).

As *Spdef^-/-^* mice showed a delayed tendency in clearing *C. rodentium* based on the tissue specific CFU values at 14-DPI, we sought to investigate whether this is due to higher load of *C. rodentium* in the crypts of these animals. To assess crypt-associated bacterial colonisation, we infected *Spdef^+/+^* an *Spdef^-/-^* mice with GFP expressing *C. rodentium* to enable fluorescent visualization of bacteria in fixed colonic whole mounts. At 14 dpi, *Spdef^-/-^* mice exhibited more frequent crypt invasion and increased epithelium-associated bacterial burden compared to *Spdef*⁺^/^⁺ littermates, in which only sparse GFP⁺ microcolonies were detected (Fig. 6L). Quantification confirmed significantly elevated mean MFI of GFP signal and a greater number of bacteria-filled crypts in *Spdef*-deficient animals (Fig. 6M).

Together, these findings highlighted the protective role of *Spdef* and the dependent icGC population in restricting early epithelial colonisation and promoting clearance of *C. rodentium* from the distal colon. In contrast, Tgm3-mediated mucin cross-linking appears dispensable in this context, despite its known significance in maintaining mucus integrity during inflammatory challenges.

### C. rodentium infection diminishes the frequency of icGCs

Given the importance of Spdef and the associated icGC population in promoting early mucosal defence and pathogen clearance, we hypothesised that *C. rodentium* may actively target this GC subset during infection.

To test this, we quantified icGCs using Atp12a (a protein that encodes the catalytic subunit of a non-gastric proton pump regulator) as a cell surface marker, as it is enriched in the icGC cluster identified by prior single-cell transcriptomic analysis (Fig. 7A) [8]. To visually confirm the spatial location of Atp12a expression, we immunostained colonic sections for Atp12a and confirmed expression in the upper crypt region, most notably within the vesicle membranes of icGCs (Fig. 7B). We therefore proceeded to infect RedMUC2^98trTg^ mice with *C. rodentium* and extracted epithelial cells across the course of the infection. Flow cytometry gating was constructed by first identifying live EpCAM⁺ mCherry-Muc2⁺ epithelial cells, followed by stratification based on Atp12a expression (Fig. 7C).

**Figure 7:**
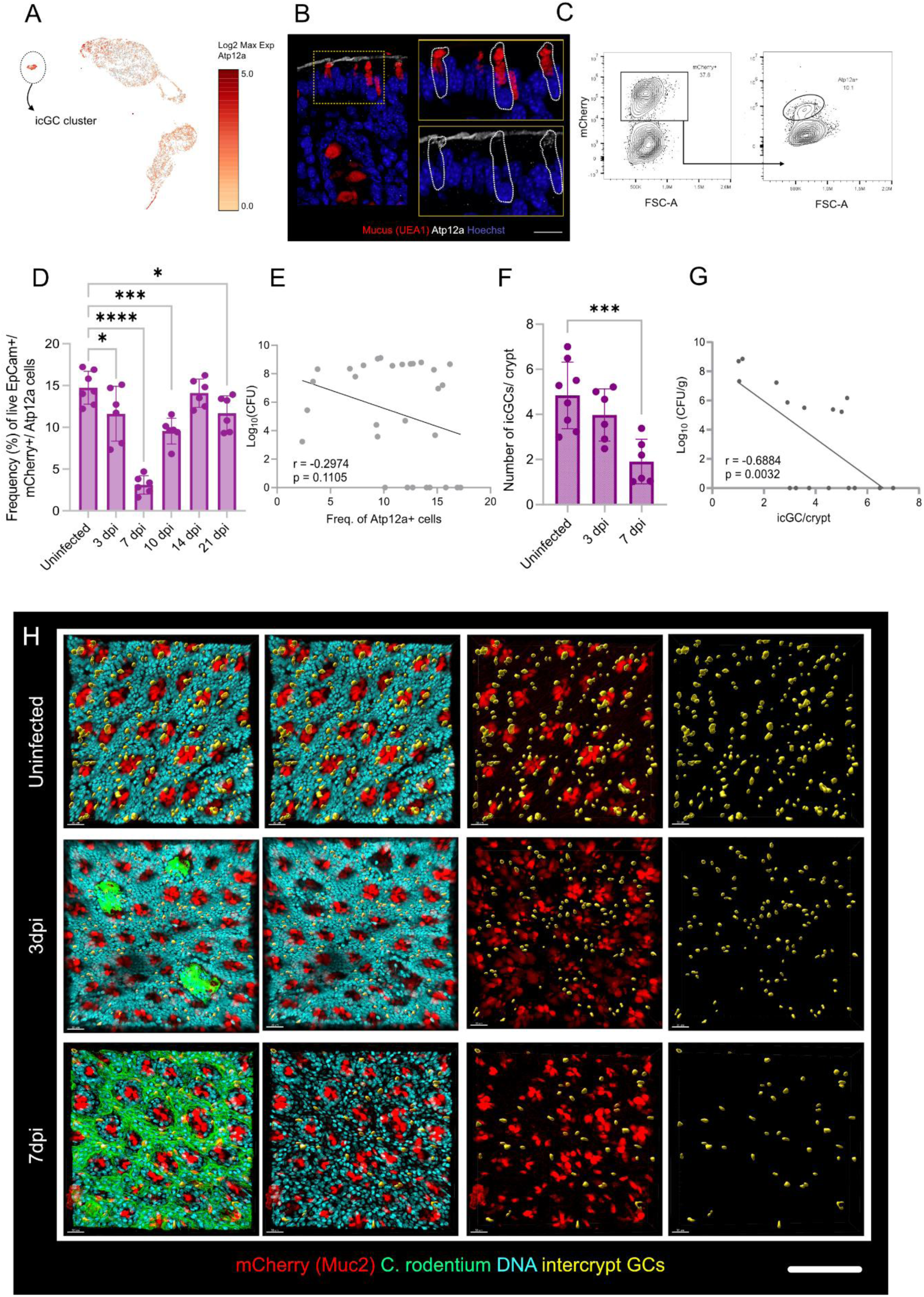
C. rodentium infection reduces the frequency of intercrypt goblet cells in the distal colon. **A)** Uniform Manifold Approximation and Projection (UMAP) of single-cell transcriptomic data showing the intercrypt goblet cell (icGC) cluster and expression of Atp12a across goblet cell populations. Colour scale indicates log₂ maximum expression of Atp12a. **B)** Representative confocal micrographs of distal colonic tissue sections stained for mucus with UEA-1 (red), Atp12a (white) and DNA with Hoechst (blue). Yellow boxes indicate regions shown at higher magnification, and white outlines indicate Atp12a-positive cells within colonic crypts. **C)** Representative flow cytometry plots showing the identification of Atp12a-positive goblet cells within the live EpCAM⁺ mCherry-Muc2⁺ epithelial cell population. **D)** Frequency of Atp12a-positive goblet cells, identified as live EpCAM⁺ mCherry⁺ Atp12a⁺ cells, among total goblet cells in uninfected and C. rodentium-infected mice at 3, 7, 10, 14 and 21 days post-infection (dpi). **E)** Correlation between the frequency of Atp12a-positive goblet cells and stool C. rodentium burden, quantified as log₁₀ CFU. Linear regression line and Pearson correlation coefficient (r) and p-value are shown. **F)** Number of icGCs per crypt quantified from distal colonic whole mounts of uninfected and C. rodentium-infected mice at 3 and 7 dpi. **G)** Correlation between the number of icGCs per crypt and stool C. rodentium burden, quantified as log₁₀ CFU. Linear regression line and Pearson correlation coefficient (r) and p-value are shown. **H)** Representative confocal micrographs of distal colonic whole mounts from uninfected and C. rodentium-infected RedMuc298trTg mice at 3 and 7 dpi. The rightmost column shows the isolated icGCs. Data represent n = 6-7 mice per group for flow cytometry analysis (D). Bar graphs show mean ± SEM with individual data points overlaid (D, F). Statistical comparisons were performed using one-way ANOVA followed by Fisher’s LSD post hoc test (D, F). Correlations were assessed using Pearson’s correlation analysis (E, G), with corresponding r and p-values indicated in each graph. Statistical significance is indicated as p < 0.05 (), p < 0.001 (**), and p < 0.0001 (****). Image scale bars represent 50 µm (B, H).

Our flow cytometric quantification revealed a significant decline in the frequency of Atp12a⁺ GCs (as a proportion of total mCherry-Muc2⁺ EpCAM⁺ goblet cells) as early as 3 dpi. This population dropped from 17% in uninfected animals to 15% at 3 dpi, significantly dropping to 4% at 7 dpi, before recovering to baseline levels (15%) by 21 dpi (Fig. 7D). Our correlation analysis between CFU burden and frequency of Atp12a+ GCs revealed a moderate inverse trend (r = –0.29, p = 0.11) possibly indicating that higher colonic *C. rodentium* loads maybe associated with a lower frequency of Atp12a⁺ GCs (Fig. 7E).

To determine whether this decline reflected diminished Atp12a expression in GCs or a true loss of icGCs, we performed whole-mount confocal imaging of colonic tissue obtained from RedMUC2^98trTg^ mice infected with GFP tagged *C. rodentium.* We stained for DNA (Hoechst), in addition to the existing mCherry-Muc2 signal to help visualization and quantification of icGCs. Our analysis confirmed a marked reduction in icGC abundance per crypt at 3 and 7 dpi compared to uninfected controls (in yellow; Fig. 7H). Quantification showed a significant decrease in total icGCs per crypt in our whole mounts: from ∼7 per crypt in uninfected mice to 5 at 3 dpi and 3 at 7 dpi (Fig. 7F). Moreover, we were able to demonstrate a significant inverse relationship between pathogen burden and icGCs/crypt, signifying a strong correlation between *C. rodentium* burden and diminishing icGC numbers (Fig. 7G).

These results demonstrate that *C. rodentium* infection leads to a transient but substantial depletion of icGCs. Given the protective role of icGCs in maintaining barrier integrity via secretion of specialized intercrypt mucus, we hypothesized that icGC loss would erode this substructure. In uninfected tissue, lectin staining delineates a continuous intercrypt mucus structure produced by icGCs (UEA-1-positive) that bridges adjacent crypt openings, while WGA highlights intact crypt-associated plumes, together forming the structure of the IML (Fig. 8A, Uninfected). At 7 dpi, corresponding to maximal icGC loss, the intercrypt substructure was markedly diminished or absent, despite preservation of WGA-positive crypt plumes indicating residual crypt goblet-cell activity (Fig. 8B, C-D). Consistent with the loss of intercrypt mucus, GFP-tagged *C. rodentium* localized into the intercrypt spaces specifically where the lattice was depleted, suggesting that disruption of the icGC-dependent substructure coincides with bacterial persistence at the intercrypt surfaces (Fig. 8E).

**Figure 8:**
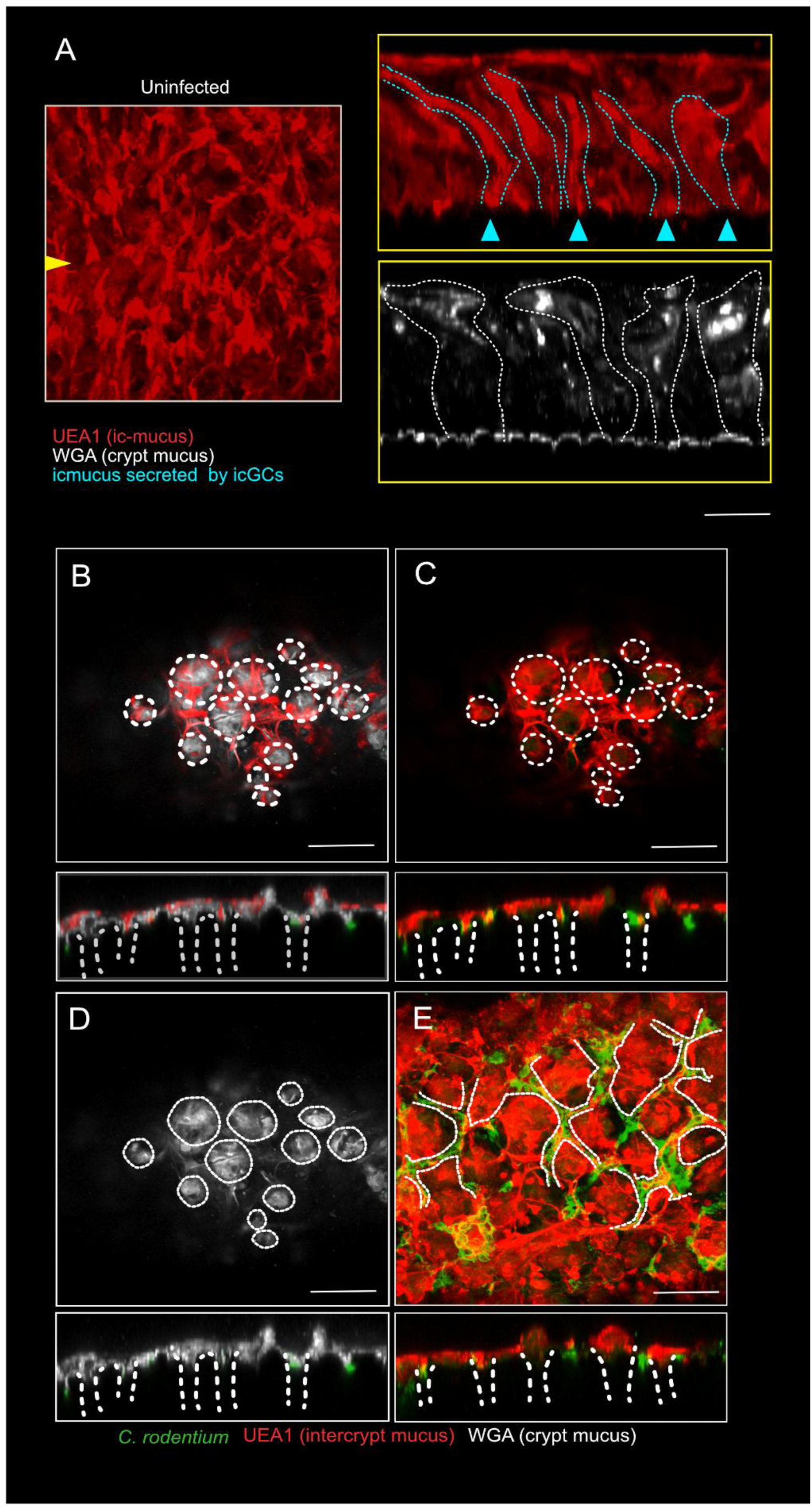
C. rodentium infection disrupts the distal colonic intercrypt mucus network. **A)** Representative confocal micrographs of uninfected distal colonic tissue stained for intercrypt mucus with UEA-1 (red) and crypt-associated mucus with WGA (white). Images show an x/y-axis view (left) and x/z-axis cross-sections (right). The yellow arrowhead indicates the location of the optical section. Cyan dashed lines and arrowheads indicate intercrypt mucus, whereas white dashed lines outline crypt-associated mucus. **B)** Representative confocal micrographs of distal colonic tissue at 7 days post-infection (dpi), showing an x/y-axis view (top) and an x/z-axis cross-section (bottom). Images show intercrypt mucus stained with UEA-1 (red), crypt-associated mucus stained with WGA (white), and GFP-expressing C. rodentium (green). White dashed lines indicate crypt openings. **C)** Representative confocal micrographs of distal colonic tissue at 7 dpi, showing intercrypt mucus stained with UEA-1 (red) and crypt-associated mucus stained with WGA (white). White dashed lines outline crypt openings in the x/y-axis view (top) and crypt structures in the x/z-axis cross-section (bottom). **D)** Representative confocal micrographs of distal colonic tissue at 7 dpi showing WGA-positive crypt-associated mucus (white). White dashed lines outline crypt openings and associated mucus structures in the x/y-axis view (top) and x/z-axis cross-section (bottom). **E)** Representative confocal micrographs of distal colonic tissue at 7 dpi showing GFP-expressing C. rodentium (green), UEA-1-positive intercrypt mucus (red), and WGA-positive crypt-associated mucus (white). White dashed lines outline crypt-associated regions in the x/y-axis view (top) and x/z-axis cross-section (bottom). Image scale bars represent 50 µm (A, x/y-axis view; B-E) and 15 µm (A, x/z-axis cross-sections).

Together, these data demonstrate that targeted loss of icGCs undermines intercrypt mucus defenses, likely driving the increased mucus permeability and epithelial susceptibility observed during *C. rodentium* infection.

### C. rodentium T3SS effector protein EspF is essential to drive intercrypt goblet cell depletion

*C. rodentium* colonizes the colonic epithelium by forming attaching and effacing (A/E) lesions, a process mediated by its type III secretion system (T3SS). This system translocates a suite of bacterial effectors including, but not limited to Tir, EspF, EspH, EspI into host epithelial cells to manipulate host signaling, cytoskeletal dynamics, and barrier integrity. Of interest EspF, has been shown to disrupt epithelial tight junctions, induce mitochondrial dysfunction, and promote epithelial cell extrusion, while the T3SS inner membrane component EscN functions as the ATPase that powers the T3SS apparatus [20, 25, 26].

To determine whether these effectors contribute to the depletion of intercrypt goblet cells (icGCs) observed during infection, we infected mice with *C. rodentium* knockout strains lacking either EspF (Δ*espF*) or EscN (Δ*escN*) and compared their effects to the WT *C. rodentium* (Fig. 9A).We quantified bacterial burden in stool and found that, as expected, mice infected with knockout strains of *C. rodentium* (both EspF and EscN) colonised comparatively weakly compared to mice infected with WT

**Figure 9:**
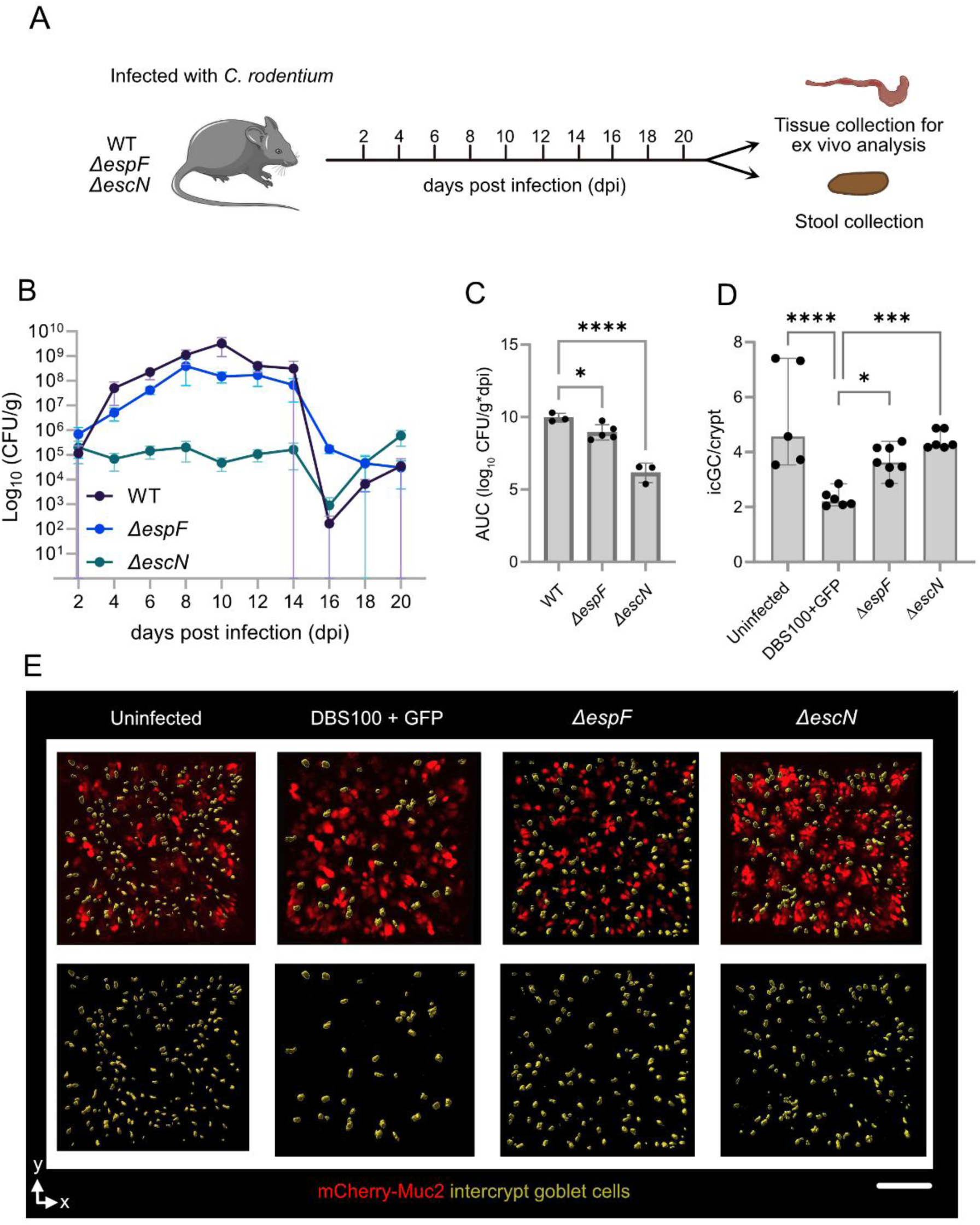
C. rodentium T3SS effector EspF contributes to intercrypt goblet cell depletion. **A)** Schematic of the experimental design using mice infected with wild-type (WT), ΔespF, or ΔescN C. rodentium. Stool samples and distal colonic tissue were collected at indicated days post-infection (dpi) for bacterial burden quantification and ex vivo mucus analysis. **B)** Stool C. rodentium burden quantified as colony-forming units per gram of stool (CFU/g) in mice infected with WT, ΔespF, or ΔescN strains over 20 dpi. **C)** Area under the curve (AUC) of log₁₀-transformed stool bacterial burden across the infection period in mice infected with WT, ΔespF, or ΔescN strains. **D)** Number of intercrypt goblet cells (icGCs) per crypt quantified from distal colonic whole mounts of uninfected mice and mice infected with WT GFP-expressing DBS100, ΔespF, or ΔescN strains at 7 dpi. **E)** Representative confocal micrographs of distal colonic whole mounts from uninfected mice and mice infected with WT GFP-expressing DBS100, ΔespF, or ΔescN strains at 7 dpi. Images show mCherry-Muc2 (red) and identified intercrypt goblet cells (yellow). The lower panels show the isolated intercrypt goblet cell signal. Data represent n = 3-5 mice per group (B, C) or n = 5-7 mice per group (D). Graphs show mean ± SEM (B) or mean ± SEM with individual data points overlaid (C, D). Statistical comparisons were performed using one-way ANOVA followed by Fisher’s LSD post hoc test (C, D). Statistical significance is indicated as p < 0.05 (), p < 0.001 (**), and p < 0.0001 (****). Image scale bar represents 50 µm (E).

*C. rodentium*. *ΔespF* reached a peak CFU in stool of 10^8^/g while *ΔescN* had a peak value of 10^6^ CFU/g stool (Fig. 9B). To further quantify differences, we compared area under the curve (AUC) values between samples infected with WT and the *C. rodentium* knockout strains. Our results demonstrate a significant reduction in AUC values among *ΔespF* and *ΔescN* infected samples when compared to samples infected with WT (Fig. 9C).

To determine if altered infection capacity was also associated with icGC depletion, we examined whole mounts from mice infected with WT, Δ*espF* or Δ*escN* by staining them for DNA (Hoechst) and mucus (UEA-1). *C. rodentium* was visualized by GFP fluorescence (WT-GFP) or by staining for LPS (*ΔespF* and *ΔescN*). Our whole mount confocal imaging further confirmed that Δ*espF* and Δ*escN* infected mice exhibited a comparatively preserved icGC frequency, in contrast to the disrupted epithelial interface and icGC depletion seen in WT infected animals (Fig. 9E). When we quantified the icGC numbers between these groups, the number of icGC per crypt in WT infection was estimated to be ∼2 per crypt by 7 dpi, whereas Δ*espF*- and Δ*escN*-infected mice retained significantly higher icGC counts (4 and 5.0 per crypt, respectively; Fig. 9D).

Thus, we show that Δ*espF* and Δ*escN* strains possess reduced icGC depletion capabilities in addition to their pattern of poor colonisation as determined by the stool CFU counts. Consequently, these findings suggest that *C. rodentium* employs EscN dependent T3SS apparatus, and potentially EspF-dependent effector delivery, as critical mediators for driving icGC loss and mucus barrier disruption during infection.

### EspF-dependent GC transcriptome remodeling potentially underlies icGC depletion

Building on our previous observation that both Δ*espF* and Δ*escN* mutants not only exhibit reduced epithelial colonization compared to WT *C. rodentium* but also fail to drive the characteristic loss of icGCs, we sought to define how EspF-mediated effector delivery reshapes goblet cell physiology. Specifically, we questioned whether EspF triggers a GC–intrinsic transcriptional program that undermines survival or stress-response pathways (rather than simply killing a pre-existing icGC subset) and thereby promotes icGC depletion.

To address this, we isolated and sorted colonic GCs and neighboring enterocytes (ECs) from reporter mice infected with either WT *C. rodentium* or the Δ*espF* mutant, then performed RNA sequencing on each population to uncover EspF-dependent transcriptional changes (Fig. 10A). We enumerated bacterial burden in stool of infected mice and, as previously observed, Δ*espF* strain colonized poorly compared to the WT C. rodentium strain (Fig. 10B). Sorting efficiency of YFP⁺ GCs was validated by flow cytometry with 13.7% YFP⁺ events in Muc2-YFP reporters versus 0% in C57Bl/6 controls (Fig. 10C). Both WT and Δ*espF* infection induced significant transcriptomic changes in ECs versus uninfected controls. However, no significant differences were detected between WT and Δ*espF* infections in this cell type (Fig. 10D). In stark contrast, goblet cells exhibited a substantial number of EspF-dependent transcriptional changes where the EspF vs WT comparison yielded a high number of differentially expressed genes within GCs (Fig. 10E), suggesting that the main effect of EspF-dependent transcriptional responses occurred in the GC compartment at this infection timepoint. In line with this, EC PCA (Fig. 10F) showed substantial overlap among uninfected, WT, and Δ*espF* samples, indicating minimal global transcriptomic differences between WT and Δ*espF* in enterocytes. However, GC PCA (Fig. 10G) revealed a moderate separation by condition, with WT and *ΔespF* forming distinct clusters from uninfected and from each other, highlighting EspF-dependent remodeling of the goblet cell transcriptome.

**Figure 10:**
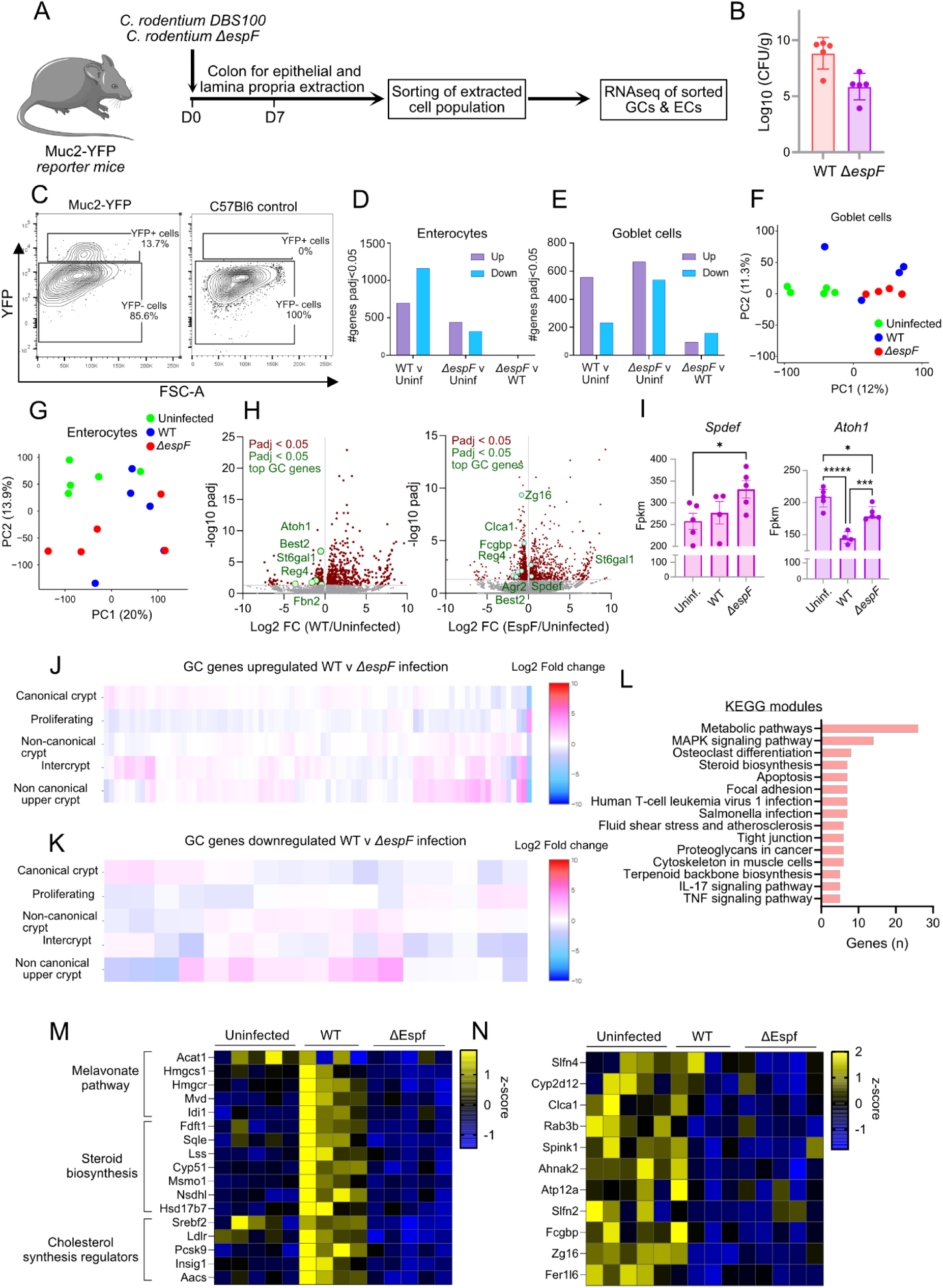
EspF-dependent goblet cell transcriptomic remodeling is associated with intercrypt goblet cell depletion. **A)** Schematic of the experimental design using Muc2-YFP reporter mice infected with wild-type (WT) or ΔespF C. rodentium. Colonic epithelial and lamina propria cells were isolated at 7 days post-infection (dpi), followed by fluorescence-activated cell sorting of goblet cells (GCs) and enterocytes (ECs) for RNA sequencing. **B)** Stool C. rodentium burden quantified as colony-forming units per gram of stool (CFU/g) in mice infected with WT or ΔespF strains at 7 dpi. **C)** Representative flow cytometry plots showing YFP fluorescence in colonic epithelial cells from Muc2-YFP reporter mice and C57BL/6 controls. YFP-positive and YFP-negative populations are indicated. **D)** Number of upregulated (purple) and downregulated (blue) differentially expressed genes (DEGs; adjusted p-value [padj] < 0.05) in ECs following WT or ΔespF infection compared with uninfected controls, and between WT and ΔespF infection. **E)** Number of upregulated (purple) and downregulated (blue) DEGs (padj < 0.05) in GCs following WT or ΔespF infection compared with uninfected controls, and between WT and ΔespF infection. **F)** Principal component analysis (PCA) of GC transcriptomic data from uninfected (green), WT-infected (blue) and ΔespF-infected (red) mice. **G)** PCA of EC transcriptomic data from uninfected (green), WT-infected (blue) and ΔespF-infected (red) mice. **H)** Volcano plots of differentially expressed genes in GCs following WT infection compared with uninfected controls (left) and ΔespF infection compared with uninfected controls (right). The x-axis indicates log₂ fold change (log₂FC), and the y-axis represents −log₁₀ adjusted p-value (padj). Differentially expressed genes are highlighted in red, with selected goblet cell-associated genes indicated in green. **I)** Expression of the goblet cell differentiation-associated transcription factors Spdef and Atoh1, quantified as fragments per kilobase of transcript per million mapped fragments (FPKM), in uninfected, WT-infected and ΔespF-infected mice. **J)** Heatmap showing the expression patterns of GC-associated genes upregulated during WT infection compared with ΔespF infection, mapped across five GC populations: canonical crypt, proliferating, non-canonical crypt, intercrypt and non-canonical upper crypt GCs. Colour scale represents log₂FC. **K)** Heatmap showing the expression patterns of GC-associated genes downregulated during WT infection compared with ΔespF infection across the same five GC populations. Colour scale represents log₂FC. **L)** Kyoto Encyclopedia of Genes and Genomes (KEGG) pathway enrichment analysis of EspF-associated differentially expressed genes in GCs. Bars indicate the number of genes associated with each enriched pathway. **M)** Heatmap showing the expression of genes involved in the mevalonate pathway, steroid biosynthesis and cholesterol synthesis regulation in GCs from uninfected, WT-infected and ΔespF-infected mice. Colour scale represents gene expression z-scores. **N)** Heatmap showing the expression of selected intercrypt goblet cell-associated genes in GCs from uninfected, WT-infected and ΔespF-infected mice. Colour scale represents gene expression z-scores. Data represent n = 5 mice per group (B) and at least three independent biological replicates for transcriptomic analyses, as indicated. Stool bacterial burden is presented as mean ± SEM (B), and gene expression measurements are shown as mean ± SEM with individual data points overlaid (I). Differential expression analysis was performed using DESeq2, with statistical significance determined by adjusted p-values (padj). Statistical significance is indicated as padj < 0.05 (*) and padj < 0.00001 (*****) in panel I.

Next, we examined GC transcriptomic changes during WT and *ΔespF* infection. This revealed that during WT infection versus uninfected controls, 19 top GC markers including *Atoh1, St6gal1* and *Reg4* were significantly downregulated (Fig. 10H, left). When comparing EspF infection to uninfected samples, 47 key GC-enriched genes were differentially regulated, notably downregulation of factors *Agr2, Clca1* and *Fcgbp* alongside a notable upregulation of *Spdef*, the transcription factor essential for icGC function (Fig. 10H, right). Consistent with this, examination of RNA expression levels confirmed elevated *Spdef* specifically in *ΔespF*-infected GCs, while *Atoh1* was suppressed in WT infection and largely restored in Δ*espF* infection (Fig.10I). These patterns indicate that Δ*espF* infection selectively shapes the GC transcriptome in ways distinct from WT infection and may confer the host with *Spdef* and *Atoh1* dependent protective responses.

To further identify which GC subsets were most affected, we overlaid these gene sets onto the single-cell GC transcriptomic dataset (Fig. 6B). Genes more highly expressed in WT versus Δ*espF* (Fig. 10J) included several with higher expression in the intercrypt and non-canonical upper-crypt clusters, including icGCs, whereas the downregulated set (Fig. 10K) was sparse in these clusters and shows minimal representation among icGCs. Together, these results indicate that EspF-dependent transcriptional changes include genes that are preferentially engaged in upper-crypt/intercrypt GC subsets, with icGCs most prominently affected.

To associate biological functions among the altered genes, we performed KEGG pathway enrichment and found pathways upregulated in WT but not Δ*espF* infection including MAPK signaling as well as steroid and terpenoid backbone biosynthesis, considered as core components of cholesterol synthesis (Fig. 10L). When visualized as a module, mevalonate/steroid pathway genes are coordinately induced in WT but not Δ*espF*, consistent with a GC program responding to sterol scarcity (Fig. 10M). This pattern suggests that WT infection drives a cholesterol-demand or depletion state in GCs, likely a membrane and ER-stress–linked response, whereas loss of EspF blunts this metabolic reprogramming. However, despite these shifts, classical icGC markers do not differ between WT and Δ*espF*, arguing against true maintenance of steady-state icGCs in Δ*espF* infected mice (Fig. 10N), suggesting that EspF may not directly deplete the icGC population.

Overall, these results indicate that EspF reshapes the goblet cell response to *C. rodentium* infection, altering GC transcriptome and inducing metabolic stress, while loss of EspF alters this reprogramming. Moreover, instead of preserving icGCs, deletion of EspF likely permits greater GC survival (potentially via upregulation of *Spdef*) and accumulation of late-stage intercrypt GCs that are transcriptionally reprogrammed yet may still support barrier integrity via intercrypt mucus production.

## Discussion

In the present study, infection rapidly thinned the inner mucus layer (IML) and increased its penetrability, while the outer mucus layer transiently expanded with visible clusters of stool-associated *C. rodentium*. Correlation analyses showed that epithelial colonization was strongly associated with IML thinning, increased mucus penetrability, and higher luminal bacterial burden, whereas tissue-associated C. rodentium levels remained broadly stable. We also observed a pronounced peak in stool-associated bacterial aggregates, corresponding to the greatest degree of luminal clustering throughout the infection time course. *Spdef* and its dependent Atp12a-expressing icGCs were critical for protection against *C. rodentium* during the early phase of infection and for driving its timely clearance. Our study shows that *C. rodentium* infection selectively depletes icGCs, creating gaps in the mucus barrier. Critically, bacterial type III effector protein EspF was required to drive icGC depletion and found to be influential in reshaping GC responses to *C. rodentium* infection by altering the GC transcriptome.

Collectively, the profound changes to mucosal landscape upon *C. rodentium* infection point to a possible sequence of events: initial epithelial colonization appears to destabilize the IML, and this barrier failure is reflected in higher mucus penetrability and luminal bacterial accumulation without a matching increase in deeper tissue invasion. The steady tissue-associated burden alongside rising luminal measures suggests a cycle in which surface attachment causes local mucus loss and increased bacterial shedding, potentially facilitating bacterial dispersal to other regions of the colon and onward transmission to new hosts. Earlier views of presence of bacteria within mucus was often associated with evidence of their active penetration or invasion of the barrier [11]. With the benefit of current understanding, it has become clear that the reporting of bacterial presence ‘inside the mucus barrier’ at 6 dpi in Bergstrom et al. likely reflected not active penetration through an intact IML, but rather the presence of *C. rodentium* being shed outward into the faecal mucus layer from the tissue as the IML becomes destabilized [11]. Moreover, our results show that the transient increase in FML thickness did not come from distal colonic GCs as our mucus measurement data pointed towards a steady *ex vivo* mucus growth rate during infection thereby indicating that this increase is possibly driven by GCs present in the regions proximal to distal colon suggesting orchestrated systemic secretory response. Consistent with this, previous studies have shown that faecal mucus is primarily composed of mucus derived from proximal colon [27]. Overall, our study shows that the functional distinction between inner and faecal layers is more than architectural and is modulated by infection that drives destabilization of the barrier [16, 27–29].

Our study also examined localized changes in GC dynamics during infection. Using flow cytometry and imaging, we found that infection induces heightened GC proliferation, with concurrent increases in Ki67⁺ GCs indicative of infection-dependent acute stress. Importantly, this proliferation occurred without changes in overall GC frequency, suggesting that the response reflects a general epithelial expansion rather than selective goblet cell turnover. In response to such stress, the host may (1) initiate programmed cell death to eliminate bacterial microcolonies, (2) mount targeted immune responses via apoptosis, or (3) shed epithelial cells to disengage bacterial attachment (8-10). While other studies have reported infection-induced modulation of Muc2 expression in the distal colon [30, 31], our results specifically shows that *C. rodentium* expands the Ki67⁺ GC compartment. This hyperplasia may transiently help maintain overall distal mucus output even as the structural integrity of the IML remains compromised.

Our study reveals a dynamic interplay between *C. rodentium* and the colonic mucus barrier, driven in part by targeted modulation of the goblet cell (GC) landscape. In this context, transcription factor Spdef emerged as a critical host determinant: Spdef-deficient mice exhibited increased susceptibility to acute infection, delayed bacterial clearance, and elevated crypt invasion, underscoring the protective role of Spdef-dependent intercrypt goblet cells (icGCs) in pathogen defense. Previous studies have associated Spdef deficiency with mucus secretory defects, absence of intercrypt mucus, and dysfunctional icGCs [8]. These defects are linked to increased DSS-induced inflammation and have also been observed in human ulcerative colitis [8].

Interestingly, although Tgm3 is active in the crypt plume mucus and contributes to its structural stability via Muc2 cross-linking, its deletion does not enhance susceptibility to *C. rodentium [7]*. This suggests that crypt plume integrity alone is insufficient for protection and further supports the idea that the intercrypt mucus produced by icGCs might probably be a more critical barrier component during early infection. In line with this, our confocal imaging revealed a marked loss of intercrypt mucus during peak infection, while crypt plume secretion remained detectable, albeit at low levels. Notably, the persistence of crypt plume mucus in the absence of intercrypt mucus may reflect a destabilized state, implying that crypt plume stability could be dependent on the surrounding intercrypt matrix. Together, these findings point to the intercrypt region as the most probable route of epithelial access for *C. rodentium* and highlight the indispensable role of icGCs in maintaining barrier integrity during infection.

Another aspect central to barrier disruption is the selective depletion of icGCs. Mechanistically, this depletion appeared to require the *C. rodentium* effector EspF and/or a functional T3SS secretion apparatus powered by EscN. Previous studies have shown that EspF disrupts tight junction proteins, compromising epithelial integrity and promoting systemic dissemination of *C. rodentium* [12]. However, our data does not support the idea that loss of EspF leads to recovery of conventional, steady-state-like icGCs. Although GCs are still present in the intercrypt regions during *ΔespF* infection, their transcriptional profile does not resemble that of homeostatic icGCs. Instead, EspF appears to actively suppress GC survival and differentiation during infection. This is reflected in EspF-dependent transcriptional remodeling: WT infection downregulates the essential GC lineage factor Atoh1, while *ΔespF* infection preserves Atoh1 and induces Spdef, a pattern consistent with improved GC survival and potential maintenance of icGC function. Mapping these changes onto a single-cell GC atlas revealed that a substantial number of EspF-sensitive transcripts are enriched in upper-crypt and intercrypt GC subsets, suggesting that EspF specifically targets these populations.

Consistent with this idea, KEGG pathway analysis revealed coordinated induction of mevalonate/steroid biosynthesis modules in WT but not *ΔespF* infection, indicating a sterol-demand/depletion state in GCs when EspF is active. This lipid-metabolic reprogramming likely compromises secretory cell fitness, reduces mucus quality, and accelerates intercrypt structure failure [13–15]. In the absence of EspF, this stress response is blunted, aligning with preserved intercrypt mucus, reduced mucus penetrability, restricted crypt access, and improved pathogen clearance. Taken together, these findings suggest that EspF contributes to barrier disruption not only by targeting epithelial junctions, but also by impairing goblet cell survival and function through sterol stress–linked transcriptional reprogramming.

In conclusion, our work sheds light on a targeted bacterial strategy to disrupt mucosal defence by depleting and functionally exhausting a specialised GC subset. It positions icGCs and their regulatory network, centred on *Spdef* as critical nodes in GC-mediated primary defence, and identifies EspF-driven GC transcriptome and sterol–ER stress remodelling as a plausible mechanism linking epithelial colonisation to intercrypt barrier collapse. These insights further highlight the important role of Spdef-dependent icGCs and secreted intercrypt mucus lattice during enteric infections.

## Materials and methods

### Experimental animals

All the mice used in experiments were bred in-house under specific pathogen free conditions with unlimited access to food and water and maintained on a 12-hour light/dark cycle. They were produced from heterozygous breeding pairs and genotyped to identify their genetic background. Littermate controls were included in all experiments. All mice of the various genotypes used in the different experiments were co-housed prior to analysis. Experimental comparisons were made only between age matched animals. *Tgm3*^-/-^ and *Spdef*^-/-^ mouse models have previously been described [9, 21]. Muc2YFP reporter animals were housed and locally used at MRBII at the Gent University Hospital (UZ Ghent). Prior to sample collection, mice were anesthetized with isoflurane and sacrificed by cervical dislocation. All experimental procedures using animals were approved by the Swedish Laboratory Animal Ethical Committee in Gothenburg.

### Citrobacter rodentium inoculum preparation and infection

A Streptomycin-resistant strain of *C. rodentium,* DBS100 (stated as WT), was used in infection experiments to quantify bacterial burden. For visualizing studies, we utilised a GFP-expressing *C. rodentium* (GFP DBS100), cultured on agar plates with 30 µg/mL Chloramphenicol for selection. In experiments involving knockout strains, we employed the Δ*espF* and Δ*escN* mutants of *C. rodentium*, generously provided by Bruce A. Vallance. These variants were resistant to 100 µg/mL Streptomycin. Infection inocula were prepared by culturing the bacteria overnight in LB broth at 37°C with agitation in a rotating incubator. Mice were gavaged with 200 mL of the inoculum (8.2 x 10^8^ to 2 x 10^9^ CFU/mL). Mice were sacrificed at specific intervals post infection, and samples were collected under sterile conditions to enumerate *C. rodentium* load at different anatomical sites. Approximately 2-3 cm segments of distal colon were excised and flushed with 5 mL of sterile PBS. Both the luminal contents and flushed tissues were collected separately. Tissue samples were homogenized in sterile PBS using an Ultra-Turrax T10 disperser (IKA), which was cleaned sequentially with 70% ethanol (twice) and sterile PBS (once) between samples. Luminal contents were homogenized in PBS using a Vortex Genie 2 shaker until thoroughly mixed. *C. rodentium* enumeration was performed by serial dilution of homogenates on MacConkey agar plates containing 100 µg/mL streptomycin or the specific antibiotic selection marker, followed by overnight incubation at 37°C for bacterial CFU quantification. For each sample, a theoretical limit of detection (LOD) was determined based on the recovery of a single colony from the lowest dilution plated. Average LOD calculated for all samples of the same type for each experiment is shown on all CFU graphs.

### Preparation of tissue for ex vivo analysis

Tissue was processed for ex vivo analysis following previously established protocols [3]. Freshly dissected tissue was flushed with ice-cold, oxygenated Krebs transport buffer (116 mM NaCl, 1.3 mM CaCl₂, 3.6 mM KCl, 1.4 mM KH₂PO₄, 23 mM NaHCO₃, and 1.2 mM MgSO₄, pH 7.4) to remove luminal contents and loose mucus. After removal of the mesenteric tissue, the colon was opened longitudinally, and the muscle layer was carefully separated by microdissection. The tissue was then mounted mucosa-side up in a custom-designed horizontal perfusion chamber. For ex vivo mucus imaging and penetrability assessments, ice-cold Krebs transport buffer was added to the chamber after staining, whereas for ex vivo mucus measurements, oxygenated Krebs-glucose buffer (10 mM glucose) and Krebs-mannitol buffer (10 mM mannitol) were added to the basolateral and apical compartments, respectively.

### Quantification of IML thickness and penetrability

To assess distal colonic IML barrier formation and function, IML thickness and penetrability to fluorescent microbeads were studied. The tissue was mounted in the chamber on ice, with the mucosal side facing up. It was first stained with Rhodamine-conjugated WGA lectin (50 µg/mL; Vector Labs) and staining of the mucus layer using 647-conjugated UEA-1 lectin (50 µg/mL; Vector Labs). The tissue was incubated on ice for 15 minutes, then rinsed with 1 mL of Krebs transport buffer. A suspension of 1-µm diameter Fluosphere crimson microbeads (1:10 dilution; Thermo Fisher) in Krebs transport buffer was applied to the apical mucosal surface. After allowing the microbeads to settle onto the mucus layer for 5 minutes, the surface was gently rinsed with 0.5 mL of Krebs transport buffer to remove any excess beads. The apical perfusion chamber was then filled with 2 mL of Krebs transport buffer, and the entire setup was transferred to an LSM700 confocal microscope system (Carl Zeiss) for imaging. Tissue and microbeads were imaged by collecting confocal z-stacks using a 20× water-immersion objective lens, 405/555 nm lasers, through the Zen acquisition software (Carl Zeiss). The resulting files were imported into Imaris (Bitplane), where fluorescent signals from WGA (marking tissue) and Fluosphere microbeads were used to generate isosurfaces. The z-axis positions of both the tissue and individual microbeads were then extracted. Mucus layer thickness was determined by calculating the average distance along the z-axis between the tissue surface and the microbeads. To assess mucus penetrability, the distribution of microbeads within the mucus layer was analysed. A frequency distribution of microbead distances from the tissue was created for each z-stack using Prism 9 (GraphPad). To compare distributions across samples, each curve was first normalized to maximum frequency values to account for differences in total bead counts in each z stack. The curves were then normalised to the position of the mucus surface (the z-position with the highest microbead frequency) to adjust for variations in mucus layer thickness, with beads located above the mucus excluded from the analysis. he area under each normalized curve was subsequently calculated using R statistical software to quantify microbead penetrability into the mucus layer across samples.

### Ex vivo analysis of IML growth rate

The secretion and processing of inner mucus layer (IML) in the distal colon were assessed by measuring the rate of mucus growth ex vivo, as described previously [3]. Tissue samples were mounted in chambers with basolateral perfusion set at 5 ml/h, and the system was kept at 37°C. To visualize mucus formation on the apical surface, 10µm black latex microbeads (Polybead; diluted 1:20). Using a stereomicroscope, both the epithelial surface and the microbeads were observed, and mucus thickness was determined by measuring the distance between them with a glass micropipette connected to a micrometer. Measurements of mucus thickness were taken every 15 minutes over a 60-minute period (from t=0 to t=60 min). Transepithelial potential difference (PD) was monitored using electrodes connected to the apical and basolateral chambers via agar bridges. The stable PD readings throughout the experiment confirmed normal ion channel functionality.

### Ex vivo FML analysis

Stool samples from *C. rodentium* infected and control mice were collected during colon dissection and kept in Krebs transport buffer. To hold the stool in place during imaging, a custom slide was prepared by shaping tac clay into a dome with a central indentation. A stool pellet was gently placed into the indentation, and a chamber was positioned over it so that the apical part of the stool is extended outside the chamber. The mucus layer encapsulating the stool was stained using a solution mix of Rhodamine-conjugated WGA lectin (50 µg/ml; Vector Labs) and 647-conjugated UEA-1 lectin (50 µg/ml; Vector Labs). After 15 minutes of staining, the chamber was rinsed with 0.5 ml of Krebs transport buffer to remove excess stain. The apical compartment of the chamber was then filled with 2 ml of Krebs buffer, and the setup was transferred to an LSM700 confocal microscope (Carl Zeiss) for imaging. Confocal z-stacks of the tissue and microbeads were acquired using a 20× water-immersion objective, 555 and 647-nm lasers, and Zen acquisition software (Carl Zeiss). The images were analysed with Imaris software, where the mucus surface was mapped to WGA signal. Measurements of the mucus volume, length, and width were obtained, and the height representing the thickness of the mucus layer covering the stool was calculated using the formula: volume = length × width × height.

### Colonic whole mounting

Colonic tissue was harvested from mice infected with *C. rodentium* GFP-DBS100 and placed in oxygenated Krebs transport buffer (pH 7.2) on ice until dissection. For dissection, the tissue was moved to a dissection tray and flushed with Krebs buffer to remove luminal contents and loose mucus. The mesentery was removed, and the colon was opened longitudinally. The transport buffer was removed, and the tissue was fixed by immersion in 4% neutral buffered formalin at room temperature for one hour. After fixation, it was rinsed with PBS and permeabilised in 0.5% Triton X-100 in PBS for 15 minutes at room temperature with shaking. Triton X-100 was then removed, and the tissue was rinsed with PBS. Staining was performed by incubating the permeabilised tissue in a solution of 1:500 Hoechst-34580 (Thermo Fisher Scientific), 50 µg/mL UEA-1 (Rhodamine), and 50 µg/mL WGA (Alexa647) in PBS for one hour at room temperature with shaking. Wherever mentioned in the experimental protocol, Phalloidin (Invitrogen, A22287) at a 1:1000 dilution was used as a marker to visualize tissue surface.

*C. rodentium* (GFP) was detected using an anti-GFP antibody (Mouse, A-11122, 1:200), and the non-GFP variant was detected using the anti-lipid A antibody (PA1-73178, 1:100) when necessary. After staining, tissues were washed twice in PBS for 15 minutes each and then mounted on slides with the mucosal surface facing up. Imaging was performed using an LSM700 confocal microscope (Zeiss).

### Epithelial cell isolation from mouse distal colon samples

A 5–6 cm segment of the distal part of the large intestine referred to as the mid and distal colon was dissected. The segment was flushed using a syringe fitted with a blunt needle containing cold 10% v/v 10x Hank’s Balanced Salt Solution without Mg²⁺ and Ca²⁺, supplemented with 1 mM HEPES buffer (HBSSwo), to clear out luminal contents. During dissection and transport, DC samples were kept in HBSSwo buffer on ice. Mesenteric fat and blood vessels were removed. To prepare for digestion, the tissue was inverted using negative pressure from a syringe with a blunt needle to grip and turn the tissue inside out, exposing the luminal surface. Both ends of the inverted colon were ligated using common thread. One end was first closed by tying, then the lumen was inflated with HBSSwo and immediately tied off at the other end. This ensured full opening of the crypts, allowing optimal access of digestive solutions to all cells, even the ones at the bottom of the crypts.

For digestion, the inflated colon was incubated in 20 mL of pre-warmed HBSSwo buffer containing 5 mM EDTA, 5% v/v inactivated fetal bovine serum, and 1 mM fresh dithiothreitol (DTT; Thermo Fisher Scientific, R0861) at 37°C with shaking at 140 rpm for 30 minutes. After this incubation, epithelial cells were detached by vigorous shaking for 30 seconds. The remaining tissue was then transferred to fresh, pre-warmed pre-digestion buffer and incubated under the same conditions for another 30 minutes. The first digestion supernatant was stored at 4°C until the second incubation ended, at which point both epithelial cell suspensions were combined. Residual tissue was discarded. The pooled suspension (∼40 mL) was centrifuged at 400 rcf for 10 minutes at 4°C, and the supernatant was discarded. The resulting cell pellet was resuspended in a solution of 1 mL Collagenase Type I (2mg/mL; Sigma Aldrich, 637958) and 4 μL of DNase I (200 U/mL; Zymo Research, E1009) and incubated at room temperature for 30 minutes with gentle pipetting every 5 minutes. Following enzymatic digestion, the cell suspension was washed with 25 mL of FACS buffer (HBSSwo supplemented with 2% FBS and 1 mM EDTA), centrifuged again at 400 rcf for 10 minutes at 4°C. The supernatent was discarded and the cells were resuspended in a desired volume of FACS buffer for downstream applications.

### Distal colon epithelial cell staining for flow cytometry

The viability dye eFluor 780 (AH Diagnostics, 65-0865-14) was diluted 1:1000 in PBS and added to the cells, followed by incubation for 30 minutes on ice in the dark. After incubation, the 96-well plate was centrifuged at 1500 rpm for 5 minutes at 4°C, and the supernatant was discarded. Depending on the nature of the target molecule and antibody type, fixation and/or permeabilization were performed. For extracellular markers, staining was performed prior to fixation, either directly or indirectly, depending on whether the antibody was pre-conjugated. For intracellular targets, three permeabilization protocols were evaluated: (1) fixation with 4% PFA for 15 minutes, followed by incubation in 0.1% saponin for 1 hour, with subsequent staining steps performed using buffers containing 0.05% saponin; (2) fixation with 4% PFA for 15 minutes, followed by permeabilization with 0.1% Triton X-100 for 5 minutes; and (3) fixation and permeabilization using the FoxP3 Transcription Factor kit (eBioscience, 00-5523-00) for 30 minutes at room temperature or overnight at 4°C.

Antibodies were diluted in 200 µL of FACS buffer at their optimized concentrations. The antibodies used included APC-conjugated anti-EpCAM (CD326; 1:1000; BD Pharmingen, 563478), anti-ATP12A N-terminal (1:100; Invitrogen, PA5-100678), and anti-Ki67 (1:1500; eBioscience, 14-5698-82). For detection of ATP12A and Ki67, an Alexa Fluor 488-conjugated donkey anti-rabbit IgG secondary antibody (1:1000; Invitrogen, A21206) was used. Antibody staining was performed for 30 minutes on ice in the dark, as previously optimized. Flow cytometry was conducted using a CytoFLEX cytometer (Beckman Coulter) and CytoExpert software. Data analysis was subsequently performed using FlowJo.

### Analysis of scRNA -seq

The previously published single-cell RNA-sequencing (scRNA-seq) dataset (GEO accession: GSE157313) was analyzed using Loupe Browser (version 6; 10X Genomics) [8]. Cells with mitochondrial gene content exceeding 20% were excluded. Additionally, cells with fewer than 200 or more than 5,500 detected genes were removed, along with those exceeding 6,500 unique gene counts in the distal colon. Immune and enteroendocrine cells were excluded based on the absence of Ptprc and Chgb expression, respectively. To minimize contamination from enterocytes or colonocytes, only cells expressing Muc2 at levels greater than 2 were retained. After filtering, 15,392 genes across 6,123 distal colon cells were included in the analysis. Dimensionality reduction was performed using Uniform Manifold Approximation and Projection (UMAP), and k-means clustering (K = 10) was applied, resulting in the identification of five goblet cell (GC) clusters. GC cluster identities were defined based on prior annotation of this dataset [8].

### RNA extraction & bulk sequencing on sorted epithelial cells

Muc2YFP mice were orally infected with *Citrobacter rodentium DBS100* (WT) or *C. rodentium ΔespF* and sacrificed at 7 days post infection. Epithelial and lamina propria cells were isolated using standard chelation and enzymatic digestion (described in previous methods), stained for EpCAM, and gated as live singlets to sort YFP+ goblet cells and EpCAM+YFP− epithelial cells on a BD FACSAria III directly into TRIzol at 4°C. RNA was extracted by TRIzol following the manufacturer’s protocol: samples were mixed and incubated for 5 min, phase-separated with chloroform (0.2 mL per 1 mL TRIzol) and centrifuged (12,000 × g, 15 min, 4°C). The aqueous phase was precipitated with isopropanol (0.5 mL per 1 mL TRIzol), pellets were washed in 75% ethanol, air-dried, and resuspended in RNase-free water with column or solution; for low-input samples, glycogen was added prior to precipitation. RNA quantity and integrity were assessed by NanoDrop. Samples were sequenced by Novogene and reads were aligned to the GRCm39 *mus musculus* reference genome. Sequencing data was analysed using DESeq2 and Benjamini-Hochberg FDR testing to determine differentially expressed genes (padj <0.05) between experimental groups [22].

### Cryosectioning and Immunohistochemistry

A small tissue fragment (∼0.25 cm) from the most distal part of the colon was collected and fixed in 4% formaldehyde (HistoLab, 02176) for 24 hours. Following fixation, the tissue was rinsed and transferred to 5 mL of 30% sucrose solution and stored at 4°C. Samples were then embedded in optimal cutting temperature (OCT) compound and stored at –20°C until they were cryosectioned. Using a CM1950 cryostat (Leica), distal colon tissues were sectioned into 6 µm slices. Sections were mounted on Superfrost Plus slides (ThermoScientific, J1800AMNZ) and stored at –20°C. Prior to staining, cryosections were permeabilized with 0.1% Triton X-100 in PBS for 10 minutes. When required, sections were stained with UEA-1 and 5 µg/mL WGA for 1 hour. In Red Muc2-Tg mice, this step was omitted since goblet cells were fluorescently labeled. Nuclear staining was performed using 5 µg/mL Hoechst for 5 minutes. Distal colon sections were mounted with ProLong Antifade reagent (Molecular Probes) and allowed to cure for 2–3 hours at room temperature in the dark.

Antigens in the sections were retreived by heating the slides in a citric acid buffer using a microwave for 3 minutes until the solution reached boiling point. The slides were then gradually cooled-first for 5 minutes at a low temperature setting, followed by 20 minutes at 4°C. Tissue sections were subsequently blocked for approximately 3 hours using a 5% goat serum blocking solution. The tissue was permeabilised by incubating the sections in 0.1% Triton X-100 for 10 minutes. Primary antibody for Ki67 was used with 1:100 concentration diluted in the blocking solution and applied to the sections for overnight incubation at 4°C in the dark. Primary antibody for Atp12a was (Anti-Atp12a, HPA039526, Sigma Aldrich) used in 1:100 diluted in blocking buffer. The following day, secondary antibody (Alexa Fluor donkey anti-rabbit IgG, 1:2000, Alexa488, Invitrogen A21206) also diluted in blocking solution, was applied for 1 hour at room temperature. If additional lectin and nuclear staining was required, a 1-hour lectin incubation followed by a 5-minute DNA stain (as described previously) was performed.

### Statistical analysis

Details of the statistical analyses performed for each experiment are provided in the corresponding figure legends. The term “n” represents the number of biological replicates (individual animals) used in each experiment. Transcriptomic visualizations were generated using Loupe Browser version 6. Statistical analyses and data visualization were conducted using GraphPad Prism version 9.4.1 (GraphPad Software). Results are presented as either mean ± standard deviation (SD) or median with range, as indicated. The statistical tests used are indicated where applicable. Significance levels are indicated as follows: * P < 0.05; ** P < 0.01; *** P < 0.0001; **** P < 0.00001; ns or not specified, not significant.

### Software used

The software used for experimental data acquisition, processing, visualization and analysis included ZEN (version 2.3; Carl Zeiss, www.zeiss.com), Imaris x64 (version 9.5.0; Oxford Instruments, imaris.oxinst.com), GraphPad Prism (version 9.4.1; GraphPad Software, www.graphpad.com), Loupe Browser (version 6; 10x Genomics, www.10xgenomics.com), ImageJ (Fiji; fiji.sc) and FlowJo (FlowJo LLC, www.flowjo.com).

## Notes

### Competing Interest Statement

The authors have declared no competing interest.

